# Coevolution with a seed bank

**DOI:** 10.1101/2023.02.08.527722

**Authors:** Daniel A. Schwartz, William R. Shoemaker, Andreea Măgălie, Joshua S. Weitz, Jay T. Lennon

## Abstract

Dormancy is as an adaptation to living in fluctuating environments. It can also influence species interactions, for example, by providing organisms with a refuge from predators and parasites. Here we test the hypothesis that dormancy generates a seed bank of protected individuals that can modify antagonistic coevolutionary dynamics. We experimentally evolved a spore-forming bacterial host along with a phage parasite that can infect active but not dormant cells. Seed banks buffered population dynamics against phage infection and retained phenotypic diversity that was otherwise lost to directional selection. By storing genetic diversity, seed banks also altered the distribution of host alleles, which contributed to dampened coevolutionary dynamics. Our findings demonstrate that dormancy generates a seed bank that can modify the eco-evolutionary outcomes of host-parasite systems.

## INTRODUCTION

Coevolution arises from reciprocal evolutionary changes between two or more species. Common among mutualists, coevolution is also an important process for understanding host-parasite dynamics (Thompson 1994). For example, antagonistic interactions tend to involve selection for defense strategies, whether they be behavioral attributes or morphological characteristics, that diminish the negative effects of a parasite on host fitness (van Houte *et al*. 2016; Amoroso 2021). In turn, parasites often evolve to overcome a host’s investment into defensive strategies, which sets the stage for an arms race (Allen *et al*. 2004; Brockhurst *et al*. 2014). However, coevolution is a complex process that is affected by factors including demographics (Papkou *et al*. 2021), mating systems (Jokela *et al*. 2009), nutrition (Larsen *et al*. 2019), productivity (Lopez Pascua *et al*. 2014), dispersal (Vogwill *et al*. 2008), and traits that may be involved in other aspects of organismal fitness (Nuismer *et al*. 2007).

One trait that may affect coevolution is dormancy. When challenged by suboptimal conditions, many organisms responsively transition into a metabolically inactive state. Organisms can also hedge their bets in noisy environments and stochastically transition between metabolic states (Lennon *et al*. 2021). These dormancy strategies create a reservoir of inactive individuals known as a “seed bank”. Although not capable of reproducing, dormant individuals enjoy reduced rates of mortality. As a result, seed banks affect the evolution and ecology of populations. For example, genetic drift is reduced with a seed bank because it increases the effective population size. In addition, dormancy retains individuals in a population that might otherwise be removed by natural selection. Taken together, seed banks buffer lineages from extinction and contribute to the maintenance of diversity within a population (Shoemaker & Lennon 2018).

Seed banks also alter species interactions in ways that may affect coevolution. For example, dormancy allows competing species to coexist via the storage effect (Cáceres 1997; Wisnoski & Lennon 2021). Similarily, seed banks modify the dynamics of antagnoistically interacting species. On the one hand, dormancy allows predators to persist when prey are scarce (McCauley *et al*. 1999). On the other hand, dormancy can provide prey and host populations with a refuge against predators and parasites (Klobutcher *et al*. 2006; Bautista *et al*. 2015; Ruf & Bieber 2022). However, in some instances, parasites have evolved to exploit dormant stages of their hosts for survival and transmission (Vizoso *et al*. 2005; Sonenshein 2006; Pagán 2022). Such findings have inspired theoretical work examining the interplay between dormancy and coevolution of host-parasite dynamics, but empirical investigations are lacking (Tellier & Brown 2009; Gulbudak & Weitz 2016; Verin & Tellier 2018).

For decades, communities of bacteria and phage have been used for testing coevolutionary theory (Brockhurst & Koskella 2013). Microbial strains can be assembled, replicated, and propagated in small volumes for hundreds to thousands of generations. Fast generation times and large population sizes allow for rapid evolution, which can be tracked through longitudinal sampling (McDonald 2019). In such studies, bacteria and phage often coexist owing in part to arms race dynamics and negative-frequency dependent selection, which has been demonstrated with infection networks and genome sequencing (Flores *et al*. 2011; Betts *et al*. 2018; Kortright *et al*. 2022). The targets of selection are typically associated with the initial step of infection where tail like-structures of a phage particle attach to receptor molecules on the surface of the bacterial host (Scanlan *et al*. 2011; Betts *et al*. 2018; Kortright *et al*. 2022). The resulting coevolutionary dynamics are dependent on the fitness costs of evolved strains as well as the maintenance of diversity, including physical or physiological refugia from phages (Schrag & Mittler 1996; Hall *et al*. 2011; Igler 2022).

Microbial systems also provide a means for testing how dormancy contributes to coevolution. One of the best understood forms of dormancy is endosporulation (Paul *et al*. 2019). When challenged by resource limitation, bacteria like *Bacillus* and *Clostridium* undergo a complex development process that transforms an actively growing vegetative cell into metabolically inert and long-lived dormant spore (Higgins & Dworkin 2012; Setlow 2016). The pathways controlling sporulation are well characterized and amenable to genetic manipulation, which can be leveraged in experimental evolution trials (Zeigler & Nicholson 2017). While sporulation confers tolerance to a broad range of environmental stressors, it may also modify interactions with phages (Butala & Dragoš 2022). For example, spores do not express receptors needed for phage attachment, which could make them resistant to infection (Chin *et al*. 1968). Nevertheless, viral parasites may be able to overcome this dormancy defense mechanism. Recent studies have demonstrated that some phages carry host-derived genes that can inhibit sporulation, eliminating any refuge that is conferred by this type of phenotypic plasticity (Schwartz *et al*. 2022, 2023).

In this study, we conducted experiments with a spore-forming bacterium (*Bacillus subtilis*) and phage parasite to test how dormancy and the resulting seed bank influences coevolutionary dynamics. After engineering a mutation in an essential gene for sporulation, we tested how seed banks affect infection rates, population dynamics, and community stability. By isolating bacteria from different time points in the experiment, we tracked the maintenance of phage resistant phenotypes in the host population in the presence and absence of a seed bank. Using pooled population sequencing, we also quantified patterns of molecular diversity and genetic signatures of coevolution.

## METHODS

### Strains and growth media

We used *Bacillus subtills* 168 Δ6 (Table S1) as the bacterial host in our experiments. This strain is free of known prophage and can form endospores (Westers *et al*. 2003). We cultured bacteria with either LB medium with low salt (5 g/L NaCl) or Difco sporulation medium (DSM; Harwood & Cutting 1990). Media were amended with chloramphenicol (5µg/ml) to which Δ6 is resistant, agar (15 g/L) for plating, and CaCl_2_ (10 mM in LB and 1mM in DSM) to facilitate phage adsorption. We used phage SPO1 as the l parasite in our experiments (Table S1). This lytic virus belongs to the Herelleviridae (Barylski *et al*. 2020) and its interactions with *B. subtilis* are well characterized (Stewart *et al*. 2009; Habusha *et al*. 2019). To amplify SPO1, we collected lysates from plate infections after flooding Petri dishes with pH 7.5 buffer (10 mM Tris, 10 mM MgSO_4_, 4 g/L NaCl, 1 mM CaCl_2_). We then cleared the phage-containing buffer from bacteria by centrifugation (7,200 Xg, 10 min) and filtration (0.2 μm).

### Phage adsorption assay

Phages initiate infection by attaching to bacterial surface molecules. We predicted that SPO1 would be unable to attach to spores owing to changes in cell surface that accompany dormancy. To test this, we conducted adsorption assays where we quantified the percentage of phage particles that attached to spores and vegetative cells over time (Kropinski 2009). We purified spores produced in an overnight culture in DSM medium by lysozyme treatment (50 µg/mL, 1hr, 37 °C), followed by SDS treatment (0.05%) and three washes in H_2_ O. To prevent germination, we resuspended purified spores in Tris-buffered saline (pH 7.5) lacking any resources required for growth. Vegetative cells were harvested from an overnight culture in LB medium, which were washed and resuspended in Tris-buffered saline (pH 7.5). To initiate an adsorption assay, we mixed 10^7^-10^8^ vegetative cells or purified spores with ∼10^4^ phages in a shaking incubator for 5 min at 37 °C before sampling. We immediately filtered samples (0.2 µm) to remove cells and adsorbed phages before measuring titer of unabsorbed phages by plaque assays. This involved double-layer plating with 0.3% agar overlays (Kauffman & Polz 2018). We measured the initial phage titer by setting up a control flask without cells. We enumerated hosts by colony plating. From phage and host abundances we calculated the adsorption rates (Kropinski 2009) and tested if they were greater than zero using a one-sided *t*-test.

### Coevolution experiment

To test how dormancy affects antagonistic coevolution, we conducted a 2 × 2 factorial experiment where we challenged bacteria against phage in the presence or absence of a seed bank (Fig. 1). To establish the + *seed bank* treatment, we cultured the wild type strain of *B. subtilis* in DSM medium, which promotes sporulation once resources are exhausted by growing cells. Because the transfer of spores into fresh medium triggers germination (see Supplementary Text), we created an external seed bank (Fig. 1). To achieve this, we harvested and washed spores twice with equal volumes of phosphate buffered saline (pH = 7.4) in a centrifuge (8,000 Xg, 5 min) to remove residual medium. Next, we heat-treated samples to kill phage and vegetative cells (80 °C, 20 min) before storing spores at 4 °C. At each transfer, spores from the external seed bank were mixed with spores from the previous transfer at a volumetric ratio of 4:1 (new:old). We added these dormant spores back to the focal population upon the next serial transfer. For the − *seed bank* treatment, we engineered a mutation in the host that disrupted its ability to make spores. Specifically, we deleted *spoIIE*, a gene that is specific to, and essential for, endospore formation, and confirmed that its deletion did not affect fitness or alter phage infection (see Supplementary Text). Within each of the seed bank treatments, we established six replicate populations that were randomly assigned to either a + *phage* or − *phage* treatment, which resulted in a total of 12 experimental units. We initiated each population by inoculating 10 mL of medium with 100 μL of an overnight culture that was started from a single colony. In the + *phage* treatment, we infected bacteria by adding 10^6^ plaque forming units (PFU) from an isogenic lysate of SPO1. We maintained the populations in 10 mL of DSM in 50 mL Erlenmeyer flasks in a shaking incubator (200 rpm) at 37 °C. Upon each serial transfer, we aliquoted 1% of the population (100 μL) to fresh medium. For populations assigned to the + *seed bank* treatment, we transferred 50 μL of an untreated population sample and 50 μL of the external seed bank to control for total inoculum size. We transferred each population every other day for 28 days for a total of 14 transfers, which amounted to approximately 90 host generations. We sampled each flask daily to quantify population sizes (see below). At each transfer (48 h), we preserved samples of the host and phage populations for assessment of phenotypic and molecular evolution (see below). Bacteria were preserved by adding glycerol (15% v/v) to a population sample prior to storage at - 80 °C. For preservation of bacterial spores, we heat treated a bacterial sample (80 °C, 20 min) before adding glycerol (15% v/v) and freezing at -80 °C. For preservation of phage lysates, 5 mL of sample was cleared by centrifugation (7200 Xg, 10 min) and the supernatant was stored at 4 °C with 0.1 mL chloroform.

**Fig. 1.**
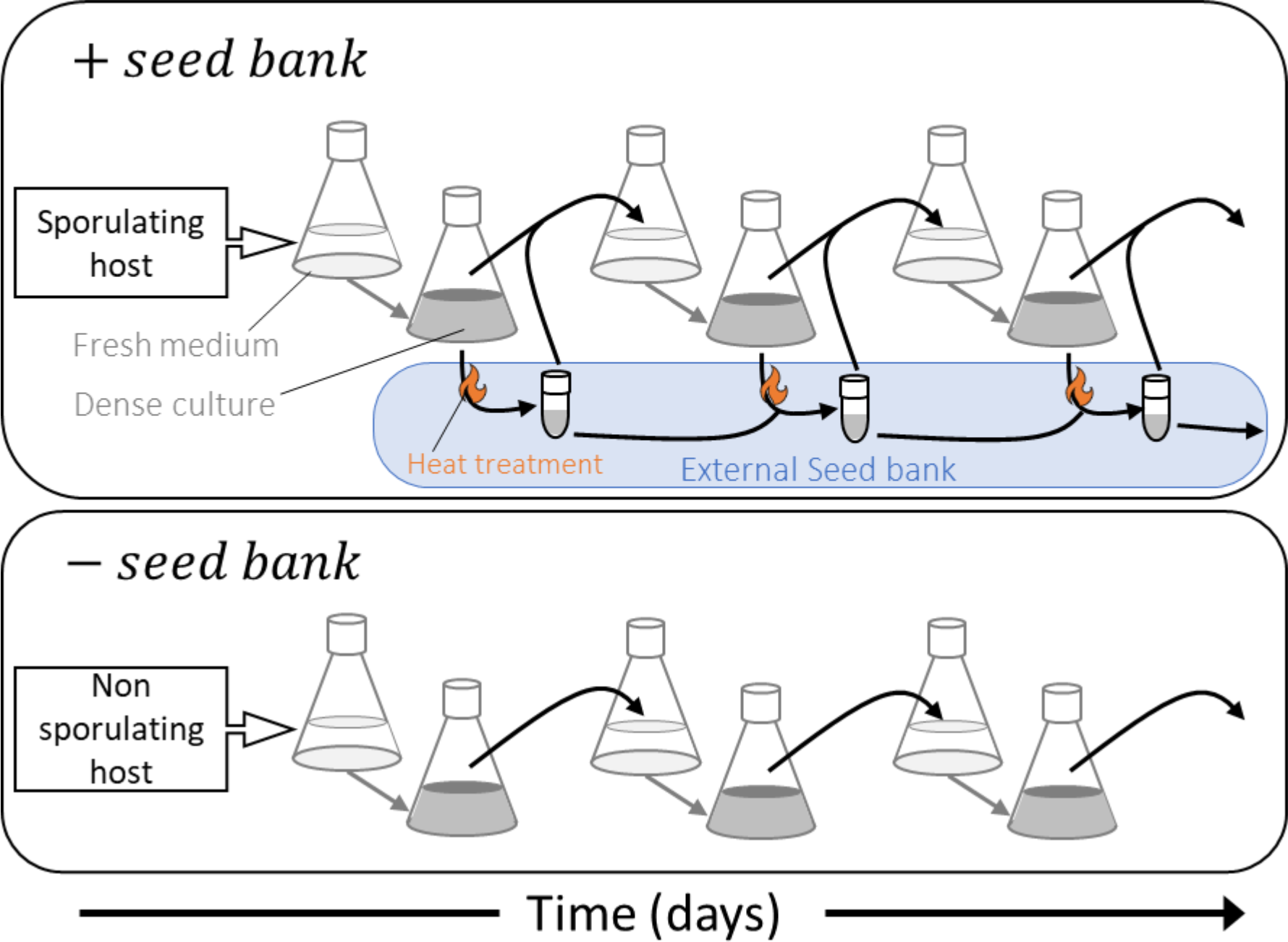
Illustration of seed bank manipulation. In the + *seed bank* treatment, wild type *Bacillus subtilis* grew rapidly in rich medium and then sporulated upon resource exhaustion before serial transfer (black arrows). In addition, we established an external seed bank to extend spore residence time. We accomplished this by heat treating samples so that only spores could survive and mixed collected spores with spores preserved from previous transfers. This spore mixture and an untreated sample were used to inoculate fresh medium and establish the next transfer. In the − *seed bank* treatment, serial transfers (black arrows) were conducted with a mutant strain of *B. subtilis* that was not capable of producing endospores in rich medium even after resource exhaustion. Each of the seed bank treatments was conducted with and without phage infection, which is not shown in the figure. See methods for further details.

### Population dynamics

We quantified bacterial densities with a flow cytometry assay that distinguished spores from vegetative cells (non-spores) based on differential uptake of the nucleic acid stain SYBR green (Karava *et al*. 2019). We quantified phage densities using a quantitative PCR (qPCR) assay with SPO1-specific primers alongside a standard curve made from a serial dilution of the ancestral phage lysate of known titer. With the resulting data, we tested for the main effects of phage treatment, seed bank treatment, and time, along with higher order interactions using repeated measures (RM)-ANOVA implemented with a linear mixed-effects model (R package nlme v3.1-149 (Pinheiro *et al*. 2020). To help meet assumptions, we transformed raw abundance data using the Box-Cox method (R package car v3.0-10, Fox & Weisberg 2018). To account for lack of independence in repeated sampling of populations over time, we included an autoregressive moving-average correlation structure (corARMA(p,q)) with parameters selected based on Akaike Information Criterion (AIC). To identify differences among treatment combinations, we conducted a *post hoc* analysis based on estimated marginal means of the RM-ANOVA model using the *emmeans* R package (v1.5.1, Russel 2020). See Supplementary Text for more detail.

### Phenotypic evolution

To evaluate how seed banks affect phenotypic evolution, we quantified host susceptibility to the ancestral phage for bacteria isolated from replicate populations over time (Fig. S1). This involved isolating bacterial clones (n = 22 per population) from samples preserved during the first four transfers of the coevolution experiment and challenging them against the ancestral phage. Clones revived from the seed bank were tested in the same manner (n = 22 per population). To characterize the susceptibility or resistance of bacteria to phage, we compared growth of clones spotted on agar plates with or without phage. Clones that could grow on both plates were scored as resistant while clones that grew only in the absence of phage were scored as susceptible. We tested for the effects of seed bank treatment and clone origin (total population vs. seed bank) on the evolution of resistance to the ancestral phage using a repeated measures ANOVA as described above.

### Molecular evolution of bacteria and phage with a seed bank

We performed pooled population sequencing to evaluate how the seed bank and phage treatments affected the molecular evolutionary dynamics of bacteria and phage populations. We extracted genomic DNA at the end of serial transfers 1, 4, 7, 10 and 14 of the coevolution experiment (see Supplementary Text). Paired-end libraries were constructed with a target minimal coverage of 100 with 2 × 38 bp reads for phage and 2 × 150 bp reads for the bacteria. Sequencing was performed using a NextSeq500 sequencer (Illumina). Mutations and their frequency were called using breseq (Deatherage and Barrick, 2014) in polymorphism mode. To focus on mutations with the largest potential effect on fitness, we restricted our analyses to nonsynonymous mutations, insertions, and deletions. Mutation frequency trajectories over time were considered only for mutations that were detected in at least three time-points.

To compare the effect of phage and seed bank treatments on the genetic diversity of bacteria, we calculated the multiplicity (*m*) of each gene in each population. Multiplicity is the excess number of mutations acquired in a gene relative to a null expectation where mutations are assumed to be randomly distributed across the genome while taking into account the size of each gene (Good *et al*. 2017). Thus, genes with high multiplicities are viewed as putative targets of positive selection. Given that few fixation events were observed, we weighed gene multiplicity by the median frequency of all mutations in that gene, excluding zeros. The multiplicity (*m*) of the *j*th gene in the *i*th population is then defined as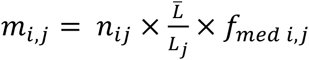, where *n*_*ij*_ is the number of mutations in gene *i* in population *j, L*_*j*_ is the number of nonsynonymous sites in the *j*th gene and 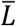 is the mean number of nonsynonymous sites among all genes, and *f*_*med i,j*_ is the median frequency of all mutations observed in gene *i* in population *j*. To account for differences in the total number of mutations acquired across populations, we normalized *m* by the sum of *m* for all genes, 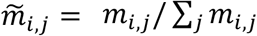. We compared the distributions of relative multiplicity across treatments using two- sample Kolmogorov-Smirnov tests with *P*-values obtained by permuting treatment labels. Last, we compared the composition of genes with putatively beneficial mutations between treatments using Principal Coordinate Analysis (PCoA) with Bray-Curtis distance. We used PERMANOVA on the top five principal coordinates (explaining >90% variation) with the *adonis2* function in vegan v2.6-2 (Oksanen *et al*. 2022) with Euclidean distance and 10,000 permutations. Results from the PERMANOVA allowed us to test for the main effects and interaction between the seed bank and phage treatment on mutated gene composition.

To determine how coevolution was affected by a seed bank, we quantified the correlation between host and phage mutations trajectories over time (McDonald *et al*. 2016). If host and phage imposed reciprocal selection on each other, we would expect there to be a strong correlation between segregating mutations of the two populations. We calculated Pearson’s correlation coefficient between pairs of mutation trajectories in corresponding host and phage populations. To minimize undue influence of zeros, trajectory pairs with less than three observations of non-zero frequencies in both host and phage populations were removed. To obtain null distributions (i.e., no coevolution), we randomly permuted time labels of observed trajectories before calculating correlation coefficients, as described above. All comparisons between distributions were performed using two-sample Kolmogorov-Smirnov tests with *P*-values obtained by permuting treatment labels. See Supplementary Text for more detail.

## RESULTS

### Phage were unable to attach to spores

Phage attached to vegetative cells at a rate of 4.63 (± 3.07) × 10^−9^ mL^-1^ min^-1^ (Fig. 2). In contrast, phages were unable to adsorb to purified dormant spores produced by the same bacterial genotype (one-sided *t*-test, *t*_*3*_ = -1.23, *P* = 0.85).

**Fig. 2.**
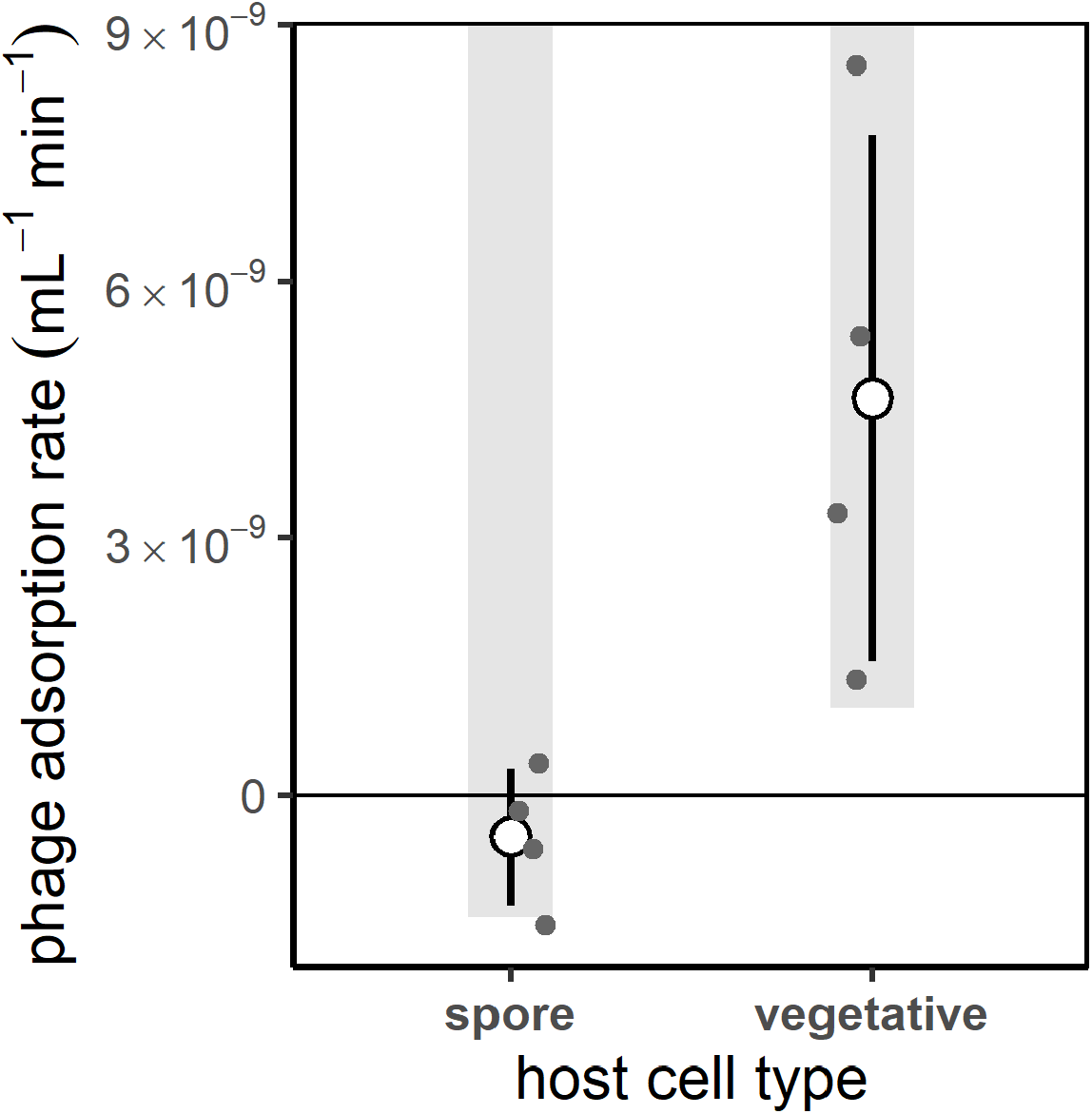
Bacterial seed banks serve as refuge from phages. Phage SPO1 cannot attach to spores of wild type *Bacillus subtilis*. Adsorption rates were calculated as the decline in free phages over 5 min when mixed with either purified spores or vegetative cells. Mean (∘) and standard deviation of four biological replicates (●) are shown. Grey bars indicate the lower limit of the 95% confidence interval.

### Seed bank altered population dynamics

Bacteria and phage coexisted for the duration of the experiment (Fig. 3). However, seed banks altered how phage affected host dynamics (RM-ANOVA; phage x seed bank x time, *F*_28, 224_ = 2.20, *P* = 0.0009). Without a seed bank, phage infection resulted in a 15-fold reduction in bacterial population size (*post hoc* comparisons based on estimated marginal means, *t*_*8*_ = 10.58, *P* < 0.0001) with minimum host densities (8.5 × 10^5^) occurring earlier in the experiment (day 5). With a seed bank, host populations, comprising both vegetative cells and spores, maintained higher densities in the face of phage infection (Fig. S2). Phage reduced population sizes by only six-fold (*post hoc* comparisons based on estimated marginal means, *t*_*8*_ = 9.68. *P* < 0.0001) with minimum host densities (3.2 × 10^7^) occurring later in the experiment (day 13). Phage-induced fluctuations in host density were stabilized by the seed bank (*post hoc* comparisons based on estimated marginal means, *t*_268_ = 2.97, *P* = 0.031). This stabilizing effect was most pronounced early in the experiment, when phages had the largest effect on host populations (Fig. S4). Without phage, seed banks had no effect on bacterial densities (*t*_8_ = 1.16, *P* = 0.28; Fig. S4), but they did increase population stability throughout the experiment (*t*_268_ = 12.50, *P* < 0.001, Fig. S4).

**Fig. 3.**
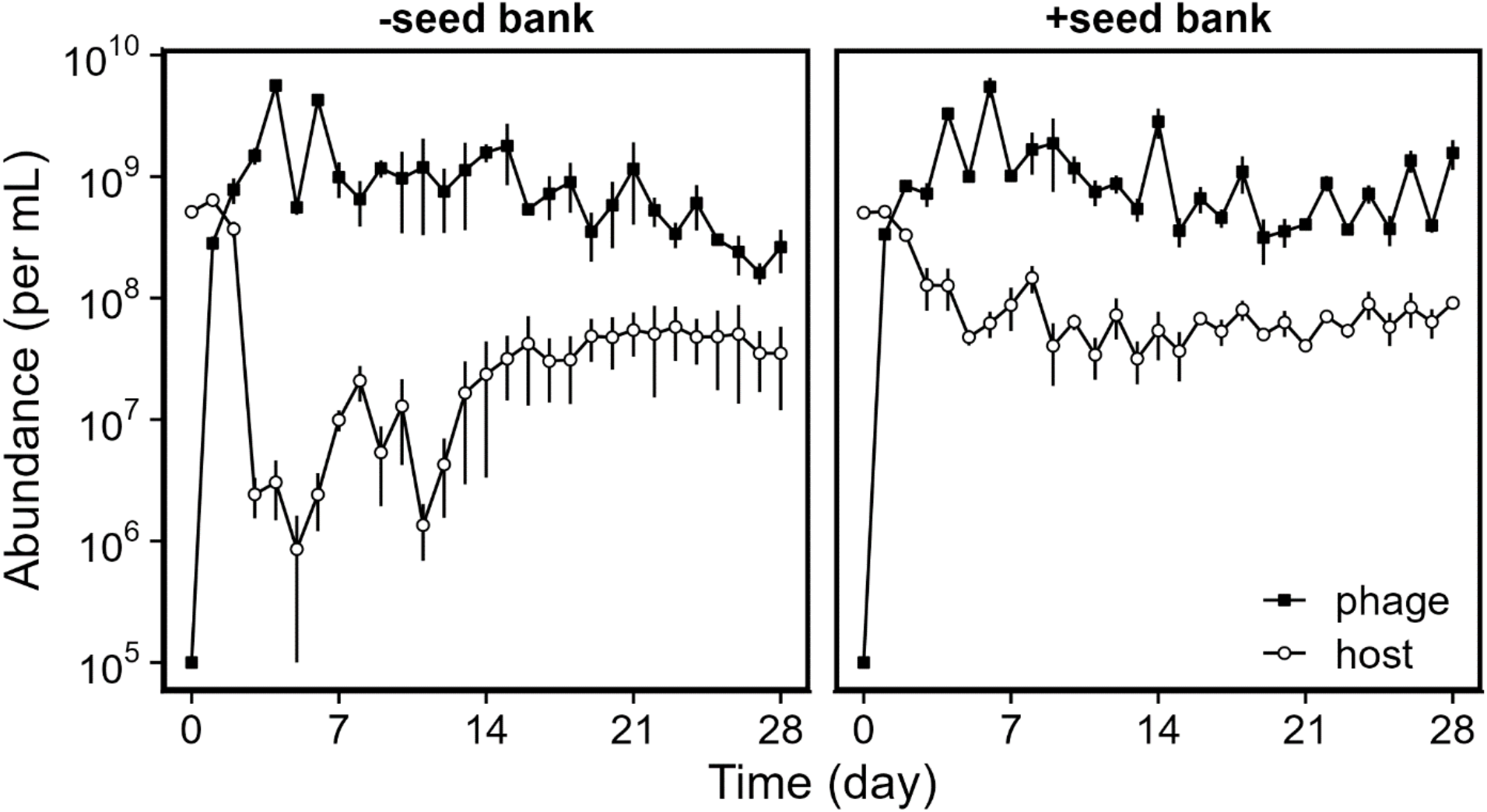
Seed banks altered host-phage population dynamics. Bacteria and phage dynamics were tracked in replicate (n = 3) populations that were propagated by serial transfer every two days. In the + *seed bank* treatment, the ancestral host could sporulate. In the −*seed bank* treatment, the host had an engineered mutation that prevented sporulation. Phage SPO1 was added to all populations at day 0. See Fig. S4 to compare population dynamics of the different host strains in the −*phage* and + *phage* treatments. Data represented as mean ± SEM.

### Seed banks retained phenotypic diversity

Without a seed bank, the ancestral phage-susceptible phenotype rapidly dropped to frequencies that were below detection limits. By the time of the first transfer, phage-resistant bacterial mutants reached nearly 100% of the population and remained so for the remainder of the experiment (Fig. 4). With a seed bank, a similar trend was observed for clones that were sampled from the total population (vegetative cells + spores) (RM-ANOVA, *F*_1, 4_ = 1.10, *P* = 0.354). However, for clones that were revived from the external seed bank, we found that the susceptible phenotype made up to 25% of the population and persisted for a longer period of time during the coevolution experiment (RM-ANOVA, *F*_1, 14_ = 12.5, *P* = 0.003).

**Fig. 4.**
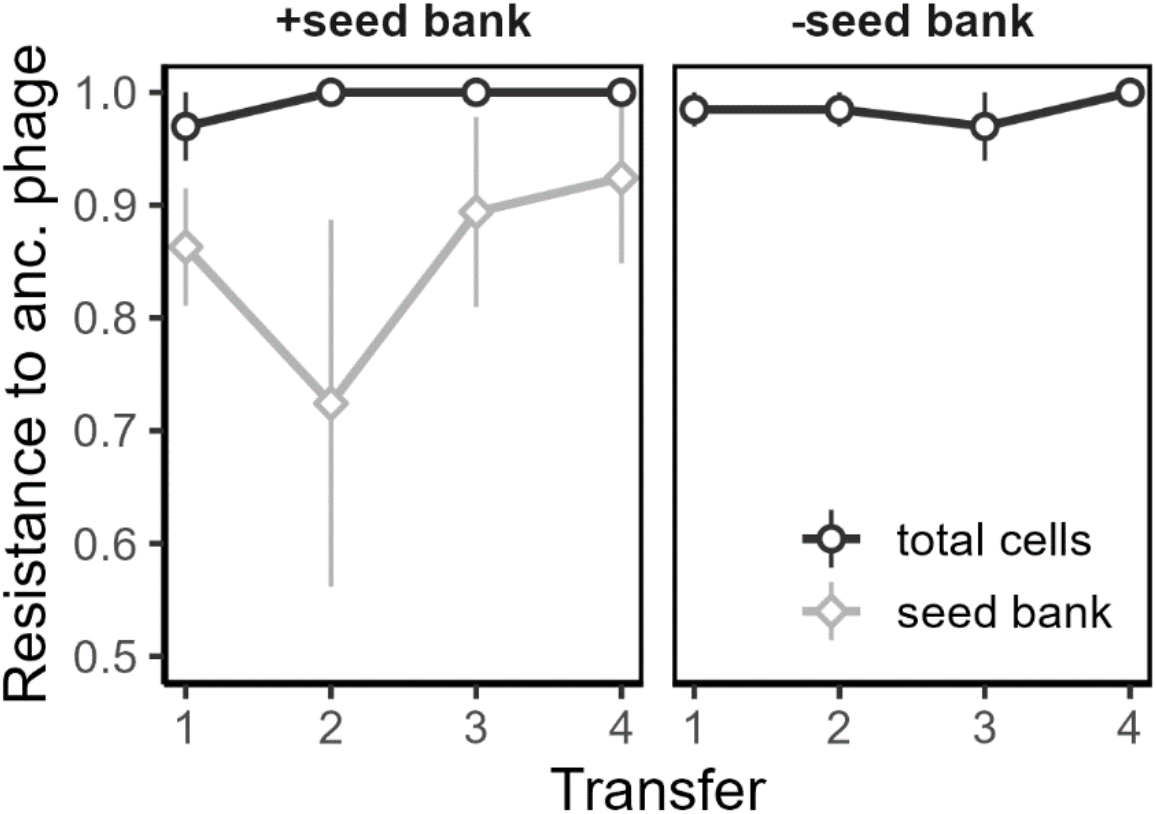
Susceptible hosts are retained in the seed bank. To characterize the evolution of resistance, clones of *Bacillus subtilis* (n = 22) were isolated from populations that had coevolved with phages and then challenged against the ancestral SPO1 phage. Total cells (vegetative + spores) were sampled from cultures prior to serial transfer. Seed bank samples represented spores that were isolated from the culture by heat treatment and mixed with spores isolated from previous treatments (see Fig. 1). Data represented as mean ± SEM of the populations (n = 3).

### Seed banks retained genetic diversity

The distribution of host genetic variation (i.e., alleles) observed in bacterial populations was significantly altered by the seed bank treatment (Fig. 5). When examining the number of non-synonymous mutations per gene weighted by their frequency in the population (i.e., multiplicity), populations evolving with a seed bank had roughly twice as many genes with mutations than those evolving without a seed bank. The additional genetic diversity from the seed bank resulted in a longer tail of multiplicity scores (Fig. 5). This pattern arose from genes in which the number of mutations, and their frequency in the population, were low, both being features that contribute to low multiplicity (Fig. 5a).

**Fig. 5.**
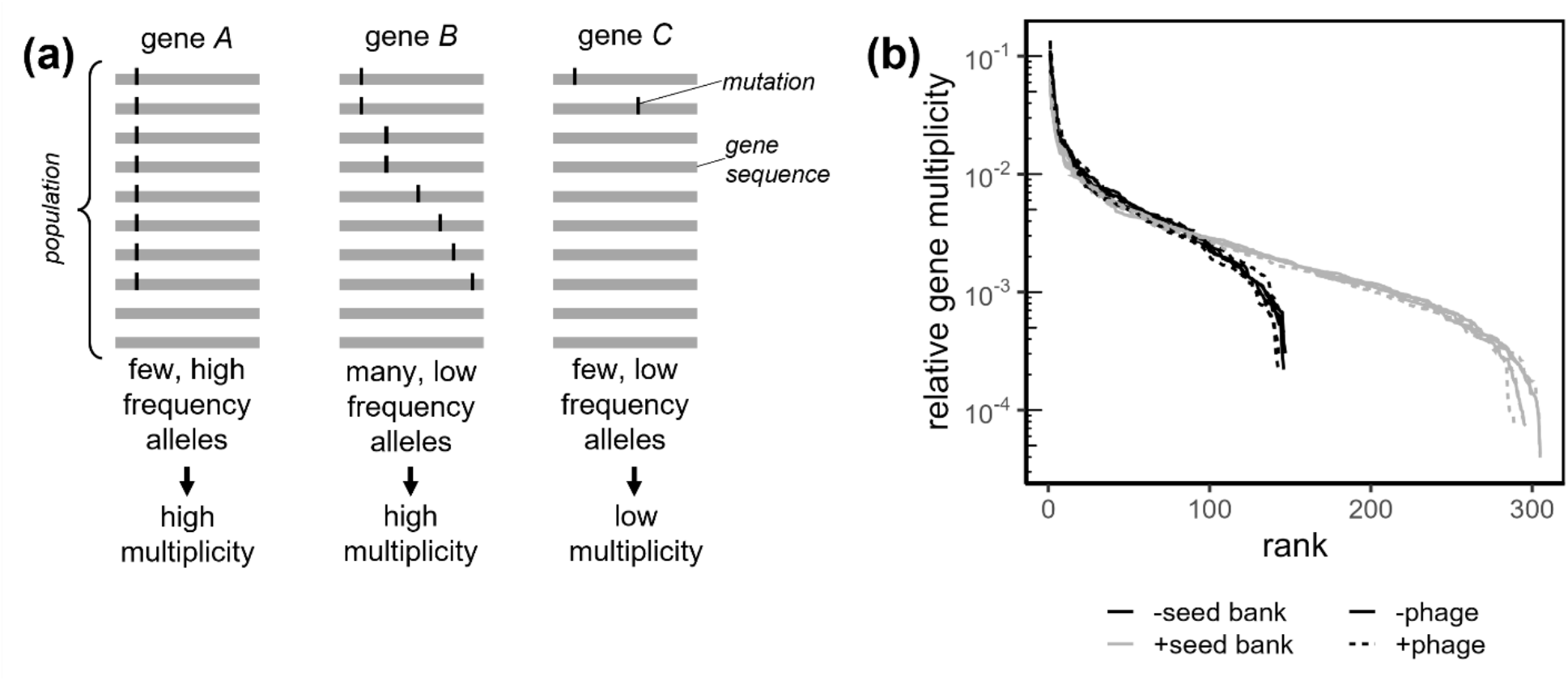
Rank-abundance distribution of mutations in host populations with and without a seed bank. **(a)** Mutations were identified by sequencing and mapped to the genes which they affect. The multiplicity of a gene reflects the excess number of mutations observed in a gene, beyond random expectation given its length, and was weighed by the frequency of those mutations in the population. Given genes of equal length (genes *A, B* and *C*) high multiplicity can arise from high mutation frequency in the population (gene A), multiple mutated sites (gene B), or a combination of the two. **(b)** For comparison among populations, we calculated the relative multiplicity, by normalizing the sum of multiplicities in each population to equal one. Each curve represents the relative gene multiplicity ranked by decreasing multiplicity values for a single population. Seed bank effects on the distribution of multiplicity were determined using a permutational Kolmogorov-Smirnov test.

The effect of the seed bank on genetic diversity was also reflected in the composition of bacterial genes with mutations. More than 70% of the allelic variation among population could be attributed to the seed bank effect (Fig. S5. PERMANOVA *F*_*1,8*_ = 65.26, *P* < 0.0001). Seed banks retained allelic variants of genes that were involved in a wide range of functions (Table S3, Supplementary Text). For example, putatively beneficial mutations were related to stress response (e.g., *fluC, yhdN, yceH*), cell wall synthesis (e.g., *dacA, ylmD*), and the regulation of gene expression (e.g., *yrdQ*). We did not observe any effect of seed banks on the distribution of allelic variants detected in phage genes (Fig. S6), or on the composition of those genes (Fig. S7).

### Genes contributing to coevolution

Phages also influenced the composition of host mutations (Figs. 6, S5, PERMANOVA *F*_*1,8*_ = 6.14, *P* = 0.022). Nearly all mutations that reached high frequencies (>0.3) in phage-infected populations were in genes involved in teichoic acid biosynthesis. Teichoic acid is a polymer found in the cell wall of Gram-positive bacteria that is used by phage SPO1 for attachment (Habusha *et al*. 2019). All replicate populations in the + *phage* treatment had high frequency mutations in at least one of four genes (*pcgA, tagD, tagF* and *gtaB*) in the teichoic acid pathway (Fig. 6, Fig. S8). Of these, *tagD* and *pcgA* mutations arose independently in separate populations, and were significantly correlated with phage infection in the ordination plot of host mutations (Table S3). Beside *pcgA* mutations, all populations with a seed bank that were infected by phage had high frequency mutations in the *sinR* repressor of biofilm formation. In the absence of phage, all populations had high frequency mutations in a single gene (*oppD*) of the *opp* oligopeptide transporter system, and four of six populations had high frequency mutations in the phosphorelay kinase *kinA* (Fig. S8). Without a seed bank, hosts had a greater number of high frequency mutations, including multiple genes of the *opp* operon and in *resE*, a sensor kinase regulating aerobic and anaerobic respiration. High frequency mutations in the phage genome were predominantly in genes encoding tail structural genes (*gp15*.*1, gp16*.*2, gp18*.*1, gp18*.*3*), irrespective of seed bank treatment (Fig. 6).

**Fig. 6.**
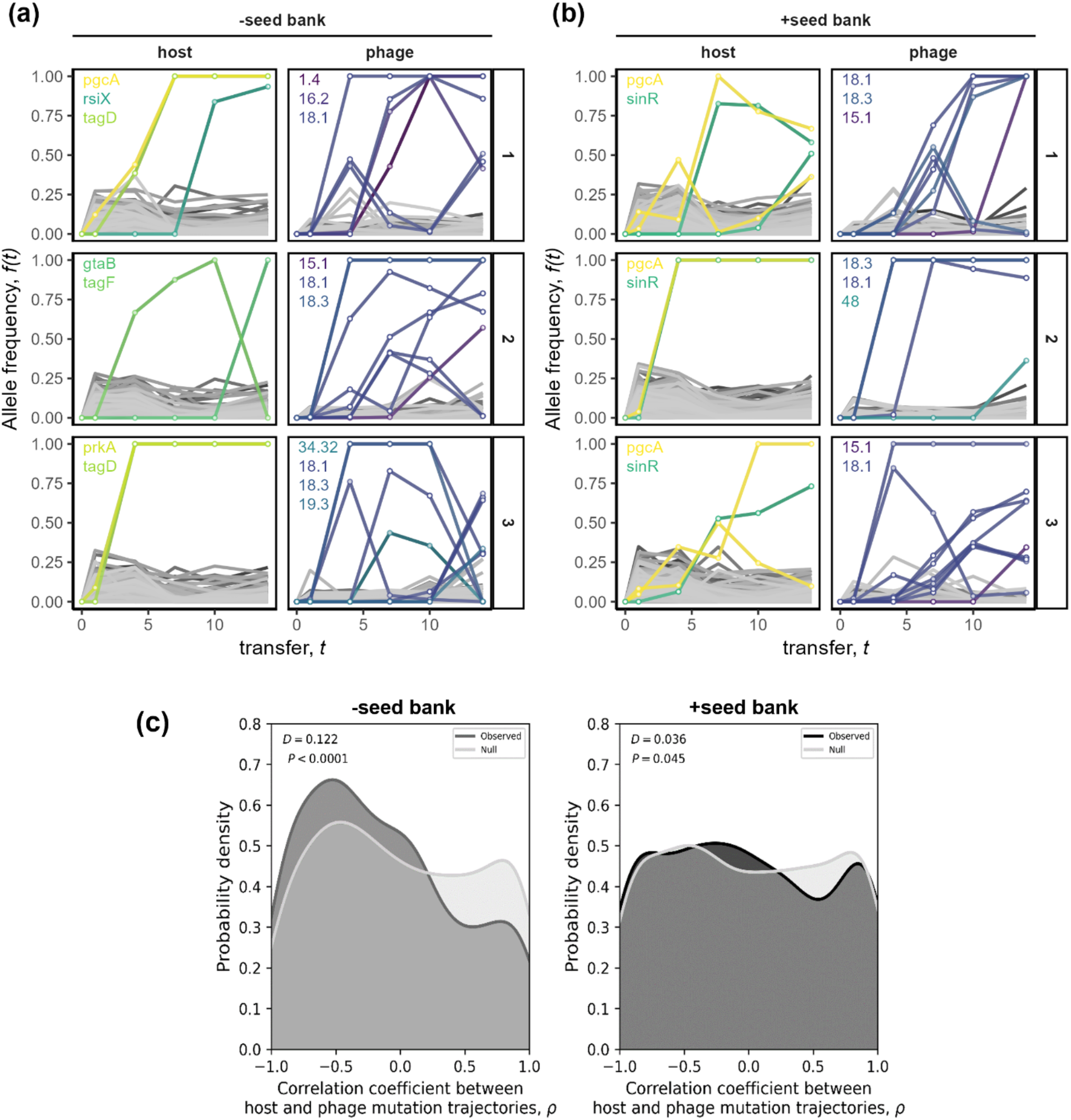
Seed banks dampened molecular coevolutionary dynamics between bacteria and phage. (**a-b)** The frequency trajectories of mutant alleles in phage-infected communities **(a)** without a seed bank and **(b)** with a seed bank. Each row shows the data for the host and phage of a single community. Non-synonymous mutations that reached a frequency > 0.3 are colored by the gene in which it occurred. The names of genes with high-frequency mutations are provided for each population. See Table S4 for details on genes. **(c)** Coevolution can lead to the coupling of mutation trajectories in host and phage populations. In the −*seed bank* treatment, the distribution of correlation coefficients between host and phage mutation trajectories skewed negative relative to a null distribution obtained by permuting time labels. In the + *seed bank* treatment, the distribution resembled that of the null with a slight overabundance of low correlation pairs, consistent with seed bank buffering of host-phage coevolutionary dynamics.

### Seed banks affect coevolutionary dynamics

Seed banks dampened the coevolutionary dynamics that typically arise between bacteria and phages (Fig. 6c). The distributions of pairwise correlations between host and phage mutation trajectories with and without a seed bank were significantly different from each other (Kolmogorov-Smirnov test, *D* = 0.151, *P* < 0.0001). Without a seed bank there was an overabundance of negative correlations compared to a null distribution obtained via permutation (Fig. 6b). With a seed bank, the observed distribution of pairwise correlations was still significantly different from the null, but its form did not skew to one side, overall being was more similar to the uniform shape of the null distribution (Fig. 6c).

## DISCUSSION

We experimentally tested how dormancy affects the coevolutionary dynamics of a bacterial host and its viral parasite. Overall, seed banks buffered population dynamics, preserved susceptible phenotypes, and retained low-frequency mutations, which led to the dampening of antagonistic coevolution. We demonstrate that a common form of microbial dormancy can reduce mortality associated with virus infection, which has important implications for understanding the persistence and spread of diseases. Because dormancy is found across the tree of life, our findings are relevant to the complexity of coevolution in other biological systems.

### Sporulation is a refuge against phage infection

Sporulation is a complex trait that is often viewed as an adaptation to fluctuating resources, but we demonstrate that is also provides resistance to phage infection. Phage SPO1 was unable to attach and infect dormant hosts (Fig. 2).

Endospores are encased in a proteinaceous coat that protects the cell wall from lytic enzymes (Klobutcher *et al*. 2006), such as those used by many phages for entry into the host cell (Fernandes & São-José 2018). Furthermore, spores do not express receptors on the cell surface that are required for attachment (Chin *et al*. 1968; Bertozzi Silva *et al*. 2016). Although transient, the spore state offers advantages compared to other forms of phage defense. It may allow the host to avoid costs associated with phage resistance mutations while also providing broad protection against multiple, or possibly all, phages (Lennon *et al*. 2007; Bull *et al*. 2014; Igler 2022). These overlooked features may help explain why spore-forming bacteria are one of the most abundant cell types on Earth (Wörmer *et al*. 2019). However, the seed bank refuge is not restricted to endosporulation. Other forms of dormancy also provide resistance to pathogens, including resting stages of bacteria, algae, plants, and metazoans (Kaplan-Levy *et al*. 2010; Brendonck & De Meester 2003; Radchuk & Borisjuk 2014; Sexton & Tocheva 2020, Pelusi *et al*. 2021).

### Seed banks alter eco-evolutionary dynamics

Phage exert strong top-down pressure on bacteria that often results in complex eco-evolutionary dynamics (Bohannan & Lenski 2000; Koskella & Brockhurst 2014). We observed such behavior in our experiment, especially in treatments where hosts lacked the ability to form endospores. Within a single transfer post infection, phage resistance swept through host populations (Fig. 3). The rapid evolution of phage resistance allowed hosts to recover and achieve densities that were comparable to non-infected populations (Fig. S4). Despite the low frequency of sensitive phenotypes, phage population sizes remained high, a pattern consistent with the invasion of host range mutants that arise owing to coevolution. In support of this, we documented an increase in the frequency of mutations associated with host receptors (teichoic acids) and phage tail components, which are hallmark targets of coevolution in bacteria-phage systems (Scanlan *et al*. 2011; Habusha *et al*. 2019; Kortright *et al*. 2022).

Seed banks significantly altered host-phage dynamics. This effect was apparent when examining host population trajectories. In the absence of a seed bank, phages reduced host densities by 15-fold with the largest effects occurring soon after infection (day 5). In the presence of a seed bank, phages only reduced host densities by six-fold and this effect was significantly delayed (day 13). Overall, this meant that host populations were more stable, most likely due to lower per-capita mortality afforded by invulnerable spores in the host population. Following each transfer, the majority of dormant individuals transitioned to active growth, a state in which they are vulnerable to phage infection. Prior to transfer the following day, resource depletion favored dormancy. In virus infected populations, this resulted in spore densities that equaled or exceeded that of vegetative cells (Fig. S2). Such findings are consistent with predictions that a refuge in the form of invulnerable prey can reduce the amplitude of fluctuations in predator-prey systems (Sih 1987; Berryman & Hawkins 2006; Pearl *et al*. 2008; Bull *et al*. 2014).

While dormancy provides benefits to the host, seed banks should also promote phage persistence, despite the evolution of host resistance to infection (Lenski 1988; Schrag & Mittler 1996). Seed banks facilitated the survival of susceptible hosts (Fig. 4), which allowed for continued phage reproduction while simultaneously experiencing losses to dilution and particle decay. Furthermore, seed banks reduce the severity of population bottlenecks, which can eliminate rare susceptible hosts and drive phage extinction during serial transfers (Hesse & Buckling 2016; Common & Westra 2019). In turn, phage reproduction supports the generation of viral diversity, which is critical for coevolution. Evolutionary models of virus-host dynamics predict that viral resistance-breaking mutants only emerge when there is continuous supply of variants that arise during viral replication (Weitz 2016). Indeed, resistance-breaking mutants are more likely to emerge in a mixed population of resistant and susceptible hosts (Benmayor *et al*. 2009; Schwartz & Lindell 2017). Overall, by stabilizing host populations and maintaining a subpopulation of sensitive hosts in a refuge, seed banks should support host-parasite coexistence and coevolution.

### Seed banks alter distribution of mutations

Consistent with theoretical expectations, our experiments revealed that seed banks maintain genetic diversity (Shoemaker & Lennon 2018; Shoemaker *et al*. 2022). The number of genes for which we detected allelic variants was roughly double in populations that had a seed bank, whether or not they were infected by phage (Fig. 5). Because our populations were initiated from single colonies, the observed host genetic diversity must have been generated *de novo* during the experiment. The increased number of variable genes in populations with a seed bank was not simply a result of having larger population size, since the pattern was also evident between seed bank treatments in non-infected populations, all of which had similar population sizes (Fig. S4). Neither can the differences be attributed to phage-accelerated diversification (Paterson *et al*. 2010), since the seed bank effect on diversity was found in infected and non-infected host populations. Rather, our results are consistent with the seed bank retaining rare alleles that would have otherwise been lost to genetic drift or negative selection. The effect on diversity recorded here at the population level is analogous to expectations of increased species richness and rarity in communities with a seed bank, which is suggested to explain long-tailed species abundance distributions observed in microbial communities (Jones and Lennon 2010). While the retention of genetic diversity is not due to phage, it has consequences for bacteria-phage coevolution. First, elevated host diversity can impede the spread of a parasite (Altermatt & Ebert 2008; King & Lively 2012). Second, prey diversity can create feedbacks that affect the dynamics and stability of predators and their prey (Steiner & Masse 2013).

### Seed banks dampen coevolution

In a coevolving bacteria-phage system, reciprocal selection should result in correlations between phage and host genotypes over time. Without a seed bank, we found that the relationship between host and phage derived alleles was skewed towards strong negative correlations (Fig. 6). With a seed bank, the correlation between segregating alleles in the phage and host populations was significantly weaker. Such decoupling likely reflects the dampening of phage selection on host variants due to the seed bank refuge. Even so, the storage of rare variants in the seed bank may ultimately enhance coevolutionary dynamics.

For example, host resistance mutations can lurk at low frequencies before rising and altering the trajectory of bacteria-phage coevolution (Gupta *et al*. 2022). When these so-called “leapfrog dynamics” arise, coevolution proceeds by an occasional replacement of the dominant host and parasite genotypes. Such dynamics are likely to be enhanced by seed bank storage. Future efforts to elucidate the net effect of seed banks on coevolution should combine phenotypic data of isolate infection networks with the genomic analysis of the host and parasite populations.

### Future directions

The use of an external seed bank has the potential to explore other ecological and evolutionary phenomena that emerge when a population has overlapping generations. Important features of a seed bank, including size and age structure, can be manipulated by altering the sample volumes and mixing ratios of the external seed bank. While tractable for use with microorganisms, in principle, the approach is amenable for use with any group of taxa where individuals can be preserved in suspended metabolic state, for example, through cryopreservation or lyophilization. Such strategies may allow for design of experiments to test theory that integrates complex life histories with demography and evolution (e.g. Yamamichi *et al*. 2019). While our experiments provide experimental control, they also bear similarity to naturally occurring seed banks where dormant individuals reside in patches that are spatially distinct from metabolically active individuals, such as plant seeds in soils or phytoplankton cysts in sediments (Moriuchi *et al*. 2000; Salazar Torres & Adámek 2013; Ellegaard & Ribeiro 2018).

Experiments like the ones described here could be expanded to explore other questions relating to the evolutionary ecology of seed banks. For example, seed banking is not limited to host populations. Parasites with dormant stages are also common (Rittershaus *et al*. 2013), and in some systems, both hosts and parasites form seed banks (Decaestecker *et al*. 2007). More work is needed to grapple with the complexity that can emerge under such conditions, but existing theory suggests that dormancy in host-parasite systems can feedback on the evolution of seed banking itself (Verin & Tellier 2018). For example, in our study system, the seed bank refuge could contribute to the maintenance of sporulation, a complex trait involving hundreds of genes. When sporulators are maintained for many generations in favorable environments, random mutations ultimately hit essential sporulation genes leading to the loss of this trait (Maughan *et al*. 2007; Shoemaker *et al*. 2022). Last, there is growing evidence that dormancy may interact with dispersal by facilitating the movement and colonization of organisms in spatially variable landscapes (Wisnoski & Lennon 2021). Experiments like the ones described here would provide a means of testing such ideas, which would be important for understanding epidemics and disease dynamics (Lennon *et al*. 2021).

### Conclusions

Dormancy is a life-history strategy that is widely distributed throughout the tree of life. It creates a seed bank, which modifies demography and diversity in ways that buffer populations against unfavorable and fluctuating environmental conditions. Seed banks also alter interactions with individuals belonging to other species, which has implications for mutualistic and antagonistic dynamics. Our study demonstrates that seed banks can serve as a refuge that stabilizes host populations when challenged by viral parasites. Protection provided by dormancy can retain genetic and phenotypic diversity of host populations with implications for understanding eco-evolutionary feedback. There is evidence, however, that parasites can exploit host dormancy behavior in ways that enhance reproductive or survivorship components of fitness (Burchard & Voelz 1972; Sastry 2013; Gabiatti *et al*. 2018; Pagán 2022). For example, phages can acquire sporulation genes, which suggests that dormancy may play an important role in bacteria-phage coevolution (Schwartz *et al*. 2022, 2023). Similar lines of investigations in other study systems will help reveal the extent to which seed banks influence the coevolutionary process.

## ACKNOWLEDEMENTS

We acknowledge technical support from Brent Lehmkuhl and Emily Long. We thank Daniel Kearns and Felix Dempwolff for strains. Research was supported by the National Science Foundation (DEB-1934554 to JTL, DAS, JSW; DBI-2022049 to JTL; and DEB-1934586 to JSW), US Army Research Office Grant (W911NF-14-1-0411, W911NF-22-1-0014, W911NF-22-S-0008 to JTL) and the National Aeronautics and Space Administration (80NSSC20K0618 to JTL). JSW was supported, in part, by the Chaires Blaise Pascal program of the Île-de-France region.

## Notes

https://github.com/LennonLab/coevolution-ts

https://github.com/LennonLab/Phage_spore_adsorption

https://github.com/LennonLab/coevo-seedbank-seq

### Competing Interest Statement

The authors have declared no competing interest.

