## Supplementary Materials for "Coevolution with a seed bank"

**Manipulating the external seed bank** — We followed a cohort of spores to test our ability to manipulate the external seed bank. We achieved this by using two *Bacillus subtilis* strains with opposing antibiotic selection markers. This allowed us to track spores of one strain when added into serially transferred populations of the other strain. We set up a serial transfer line of *B. subtilis* 3610 DK7505 (Table S1) in DSM + tetracycline (Tet, 10µg/ml). This strain is resistant to Tet and sensitive to chloramphenicol (Cm). It also constitutively expresses β-galactosidase, so it forms blue colonies on plates supplemented with X-gal (80 µg/mL). After a single growth cycle (24 h), we transferred 1% of the DK7505 population to each of two culture flasks with fresh media. At this time, we inoculated each flask with an equal volume of Δ6 spores, obtained by heat treating a sample from a stationary phase (24 h) culture of *B. subtilis* 168 Δ6 grown in DSM + Cm. Strain Δ6 is resistant to Cm, but sensitive to Tet, so could not grow in the DK7505 culture media supplemented with Tet. We then continued to serially transfer the DK7505 daily, with or without an external seed bank, as described in the Methods section of the main text. At each transfer we measured the abundance of Δ6 spores remaining in the seed bank. For this we plated seed bank samples (after heat treatment) on LB plates with chloramphenicol and X-gal (80 µg/mL). The blue/white colony phenotype allowed us to verify that there was no cross resistance of DK7505 to Cm.

In the simple transfer regime, without an external seed bank, the spiked spores decayed rapidly to extinction, within four transfers (Fig. S9). The decay rate was even greater than what would be expected by the serial dilution rate, likely due to the loss of some spores by germination in an environment unfavorable to growth (tetracycline). In contrast, application of the external seed bank treatment reduced the rate at which spores were lost from the seed bank, matching the dilution rate used in the external seed banks (1:5). Based on this we conclude that the external mixing of seed banks allows extended survival of spores in the seed bank. In this specific case the treatment extended the survival from three to ten transfers, though these numbers may vary as a function of the seed bank size.

**Deletion of *spoIIE* does not affect bacterial growth or phage susceptibility** — In the $-\,seed\ bank$ treatment of our coevolution experiment, we used a non-sporulating mutant of *B. subtilis* Δ6. To facilitate comparisons across seed bank treatments, we sought to minimize any fitness differences other than that of sporulation between host strains. To this end we deleted from *B. subtilis* Δ6 the gene *spoIIE* that is i) essential for sporulation ii) specific only to sporulation and iii) is required for the commitment to sporulation (Kuchina *et al.* 2011), so that a mutant would not be "stuck" in an early sporulation stage. The *spoIIE* gene has a dual role during sporulation initiation, being required for both asymmetric cell division and activation of spore specific gene expression during sporulation initiation (Barak & Youngman 1996). We constructed this mutant by transferring an erythromycin resistance cassette (*erm*) from a donor *spoIIE*-deleted strain (BKE00640; Koo et al. 2017) to a Δ6 recipient using generalized transduction with phage SPP1 (Yasbin & Young 1974). To eliminate fitness costs associated with antibiotic resistance, we removed the *erm* gene by transforming the strain with plasmid pDR244 (Koo *et al.* 2017). We confirmed the marker-less deletion of *spoIIE* by PCR amplification and Sanger sequencing with primers flanking the *spoIIE* locus (oDAS11 + oDAS12, see Table S2) and tested for the loss of sporulation by heat susceptibility of overnight DSM cultures. We confirmed that the *spoIIE* deletion mutant does not affect growth and phage susceptibility (see below).

We compared growth parameters of the Δ6 Δ*spoIIE* strain with its Δ6 ancestor in the sporulation media (DSM) used in the serial transfer experiment (Fig. S10). There was no difference in lag time or growth rate between Δ6 and Δ6 Δ*spoIIE* strains when cultured in DSM. As expected, there was a subtle difference in stationary phase, which is when the Δ6 strain is expected to transition into sporulation. Specifically, we found that the sporulator Δ6 strain reached a slightly higher maximum in optical density (mean $OD_{max:}$ Δ6 = 0.650, Δ*spoIIE* = 0.628; $t_{18.4} = -3.9$, $P = 0.001$).

Deletion of *spoIIE* did not affect infection by phage SPO1. The efficiency of plating was the same in plaque assays with Δ6 and Δ6ΔspoIIE as hosts ($t_{2.23} = 2.12$, $P = 0.155$). Likewise, there was no difference in virulence of phage SPO1 on the two strains of host (see Fig. S11).

**Population dynamics** — For quantifying bacteria, in triplicate, we diluted each sample in TE buffer (pH 8) and then fixed the cells in 0.5% glutaraldehyde for 15 min at 4 °C. We stained the fixed samples with SYBR green (20,000x dilution of commercial stock, Lonza) for 10 min at room temperature in the dark. We then enumerated spores and vegetative cells using a volumetric NovoCyte 2000R flow cytometer (Acea; ex 488 nm, em 530/30 nm) and an automatic gating pipeline (Karava *et al.* 2019). For quantifying phages, we used quantitative PCR (qPCR). After removing bacteria from samples via centrifugation (16,000 Xg, 2 min), we used the resulting supernatants as templates (2 µL/reaction) in triplicate 20 µL reactions consisting of iQ SYBR green supermix (Bio-rad) with 300 nM of SPO1-specific primers (qSPO1_MCP6.1_F+R, see Table S2). We ran the qPCR assays in a Mastercycler EP RealPlex thermocycler (Eppendorf) with a standard curve made from a serial dilution of the ancestral phage lysate of known titter. The PCR conditions included an initial capsid denaturation step (95 °C, 10 min) followed by 40 amplification cycles (95 °C, 15 s; 57 °C, 15 s; 72 °C, 30 s). We determined threshold cycles (Ct) using the RealPlex software with the CalQPlex algorithm, using automatic baseline and drift correction, while checking for primer dimer using melting curve analysis.

**Sequencing of bacteria and phage populations** — For sequencing of bacterial populations, we revived bacteria from cryopreserved samples taken before serial transfer (see main text methods). For revival we first washed ~10 µL of frozen samples in 1 mL of fresh DSM (8,000 Xg, 5 min) to avoid growth inhibition by glycerol (Atolia *et al.* 2020). Washed cells were grown overnight in 3 mL DSM + Cm at 37 °C with shaking (200 RPM) before we collected cells by centrifugation of 2 mL culture samples (16,000 Xg, 2 min). For gDNA extraction we treated culture pellets with lysozyme (7 mg/mL) in PowerBead solution (Qiagen) for 1 h at 37 °C. To break open cells and spores we vortexed the samples with 0.5 mm glass beads for 40 min. We then extracted gDNA using Qiagen DNeasy UltraClean Microbial Kits followed by cleanup using the Genomic DNA Clean & Concentrator-25 kit (Zymo).

For sequencing of phage populations, we removed bacteria from samples of phage-treated populations taken before serial transfer via centrifugation (16,000 Xg, 2 min) and collected the supernatants. To remove free nucleic acids, we treated supernatants for 30 min with 1.2 µg/mL DNase I and 40 µg/mL RNase at 37 °C. Next, we precipitated phages with 40 mM of $ZnCl_2$ and froze the pellets (-20 °C) until further extraction (Santos 1991). Extracted DNA was cleaned using the Genomic DNA Clean & Concentrator-10 kit (Zymo).

Sequencing libraries were constructed with Nextera XT DNA library preparation kit (Illumina) and sequenced on a NextSeq500 sequencer (Illumina). Paired-end sequencing was done as 2 x 38bp for phage populations and 2 x 150bp for host populations with a target minimal coverage of 100x. Library construction and sequencing were carried out at the Center for Genomics and Bioinformatics at Indiana University. Sequencing data are available on NCBI (Bioproject ID = PRJNA932315).

**Sequence data analysis** — Read quality of sequences was inspected using FastQC (Andrews 2010) and MultiQC (Ewels *et al.* 2016). PCR duplicates were removed using FastUniq (v1.1, Xu et al. 2012). Ancestral genomes were generated by mapping ancestor sequencing data to reference GenBank genome (host: NZ_CP015975, phage: NC_011421) using breseq in clonal mode (v0.36.1, Deatherage and Barrick 2014). To identify mutations in pooled population sequence data we mapped the reads to the appropriate ancestral genome using breseq in polymorphism mode. To identify the temporal frequency trajectories of *de novo* mutations, we applied a previously developed breseq protocol (Good *et al.* 2017) wherein the gdtools command (part of breseq) was used to consolidate mutations identified by breseq across all timepoints for each population. The merged list was used as an input for a second round of breseq, allowing for coverage values and frequency estimates to be obtained for each mutation for all timepoints within a given population.

**Functional gene categories in mutated genes** — To test if any functional genomic categories were enriched in mutated genes, we tested if gene categories proportions differed from their proportions in the entire genome. We used the first level of functional gene categories assigned by Subtiwiki (Pedreira *et al.* 2022), namely `Cellular processes`, `Metabolism`, `Information processing`, `Lifestyles`, `Prophages and mobile genetic elements`, and `Groups of genes`. For genes assigned to multiple categories, we split the count evenly between the categories so that the total number of genes was preserved. However, if `Groups of genes` was one of multiple categories, that assignment was dropped, as this category is a heterogenous catch-all category. The category proportions of mutated genes for each sample were divided into the high-ranking genes (rank ≥ 150 in Fig. 5B) and low-ranking genes. Each group of genes in each sample was tested for deviation from the whole genome category proportions using a goodness-of-fit Chi-squared test. Following multiple testing correction (Benjamini-Hochberg), all gene groups were found to be no different than the whole genome ($P > 0.1$, Table S5). However, the low ranking genes for populations with a seed bank stood out as hovering around significance, prior to multiple testing correction ($0.01 < P < 0.08$), while all high ranking genes were clearly no different than the whole genome ($0.49 < P < 0.98$). To test if any specific category was driving this trend in low-ranking genes, we used a binomial test to evaluate whether the proportion of genes in each category in each sample deviated from the whole genome proportion. Following correction for multiple testing (Benjamini-Hochberg), no single category came out as significant, suggesting there is no consistent category in low-ranking mutated genes that was enriched.

### SUPLLEMENTARY FIGURES

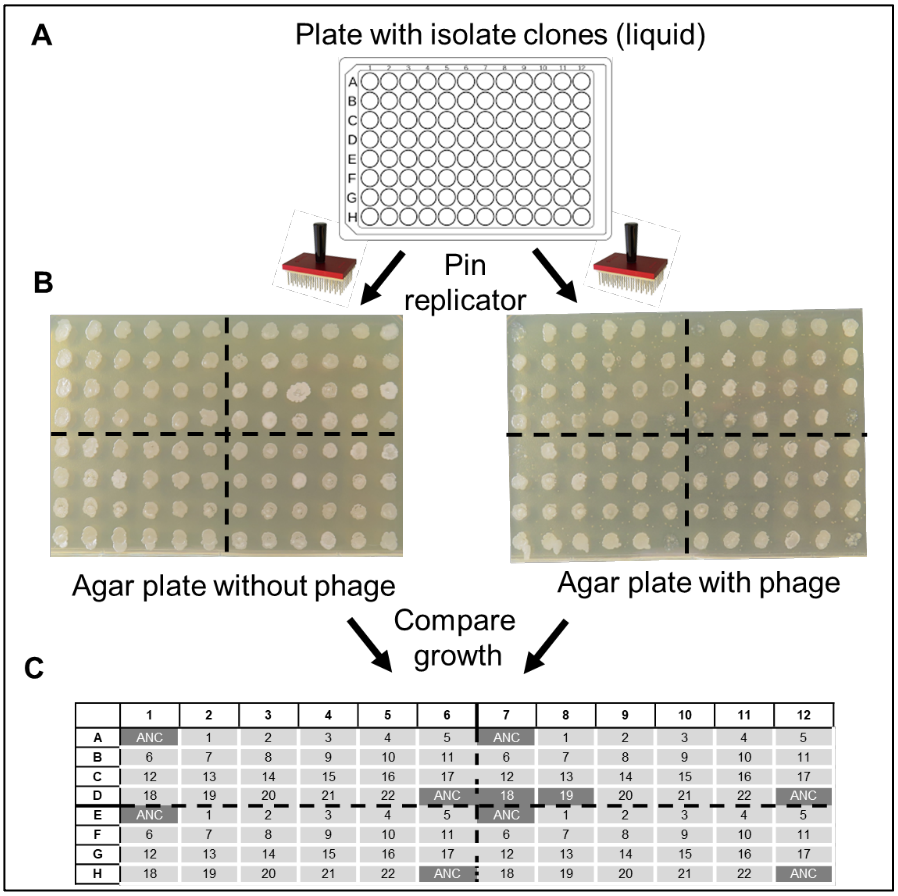


**Fig. S1**. Overview of how clone resistance was characterized using replica plating. (A) Isolated colonies were grown in liquid media (DSM) in a 96-well plate before (B) transferring culture samples using a pin replicator onto a single-well plate (OmniTray, Nunc) containing 1.5% agar DSM agar overlayed with 0.3% agar DSM with or without the ancestral phage. After overnight incubation (as shown in B) the plates were scored for growth at each spot in the array. For scoring, we only considered spots where growth was seen in the absence of phage, scoring them as resistant if they also grew in the presence of phage (light grey in C), and susceptible if not (dark grey in C). In each plate we included eight scattered wells with the susceptible ancestral host (“ANC”) to verify the distribution of phage in the entire plate. In the example shown, all clones are resistant except for #18 and #19 of the top-right group, as is the ancestor in eight spots.

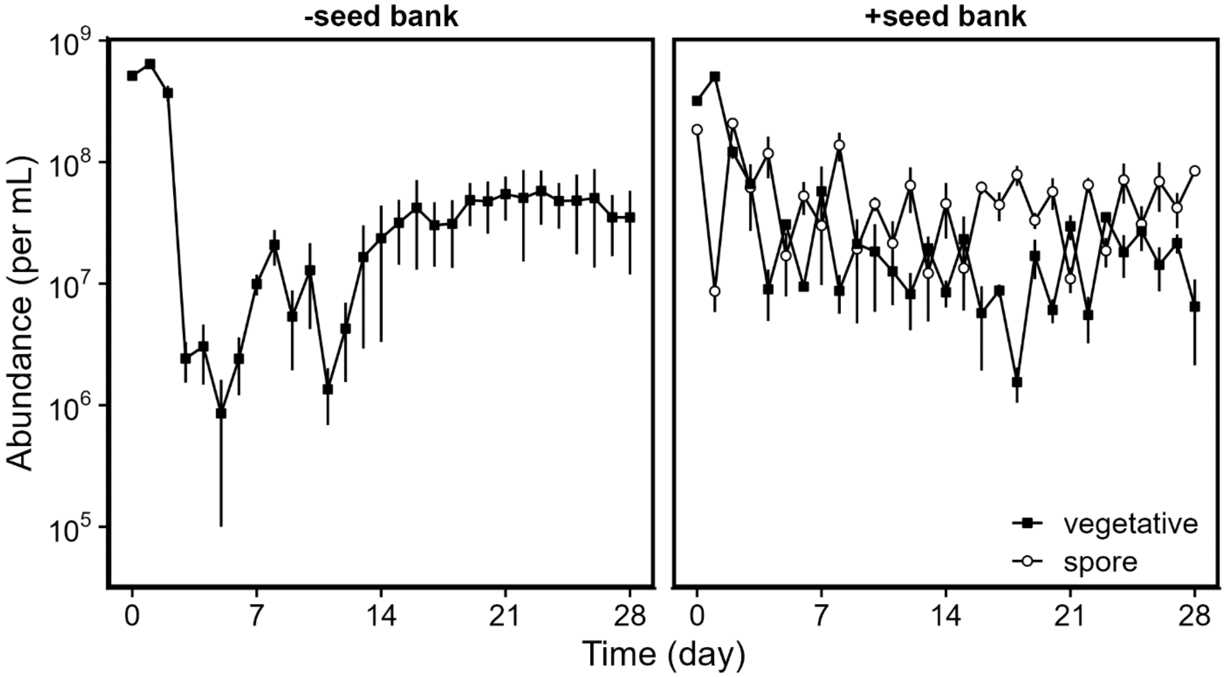


**Fig S2**. The abundance of active vegetative cells and dormant spores remained high despite being challenged by phage infection. Spore-forming ($+seed\ bank$) or non-spore-forming ($-seed\ bank$) *Bacillus subtilis* were tracked in replicate (n = 3) populations that were propagated by serial transfer every two days in sporulation media (DSM). Phage SPO1 was added to the populations at day 0. Data represent mean ± SEM.

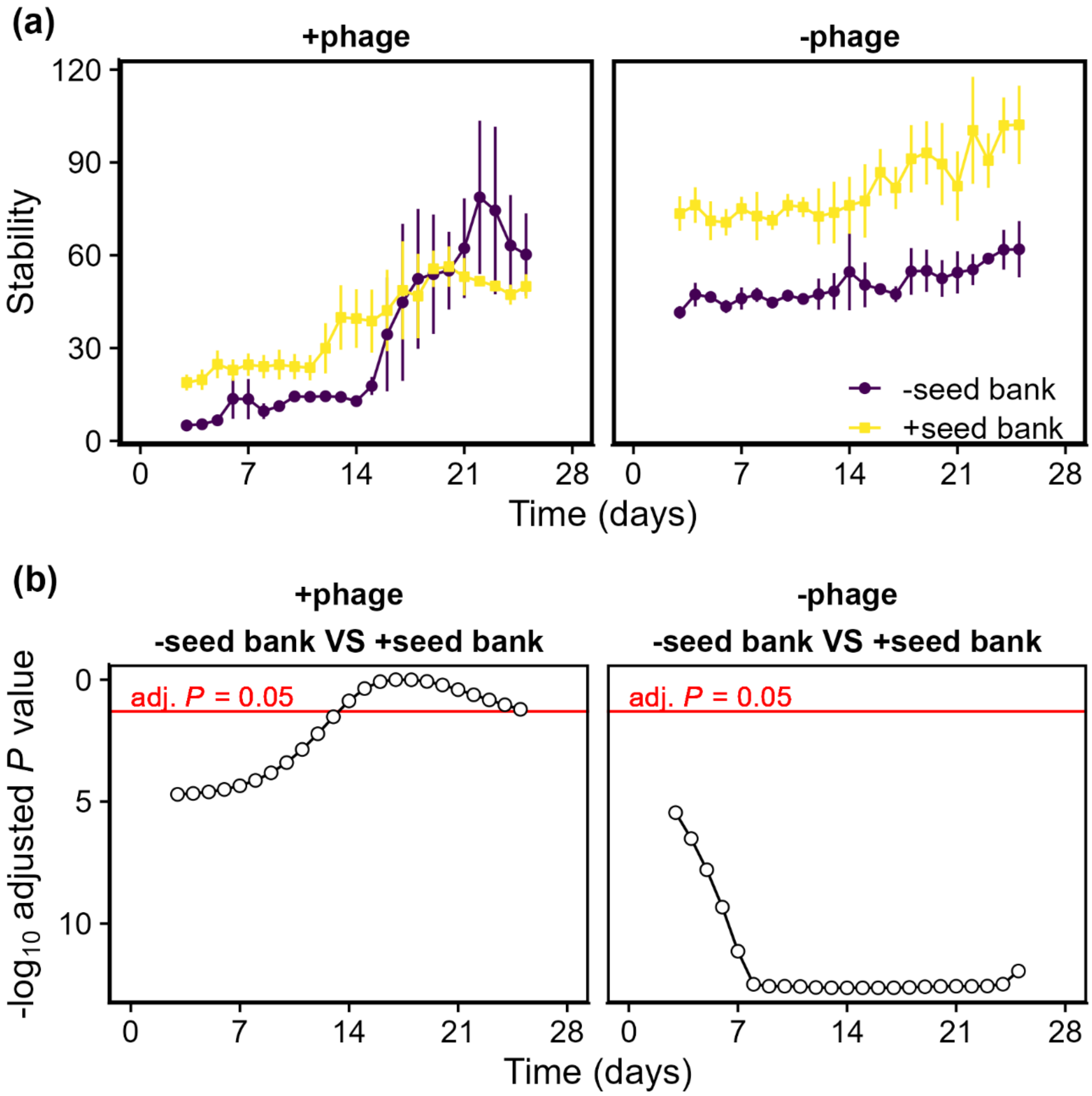


**Fig. S3.** Seed banks stabilized *Bacillus subtilis* populations. Stability was calculated as the inverse of the coefficient of variation (i.e., stability = mean/standard deviation) using $log_{10}$-transformed cell abundance data. **(a)** Each point on the plot represents stability calculated for the seven-day period surrounding that time point. For each treatment combination, replicate populations (n = 3) are represented as mean ± SEM. **(b)** *P*-values from tests comparing seed bank treatments in each time point in (a). Test results from a *post hoc* analysis based on estimated marginal means of an ANOVA model using the *emmeans* R package (v1.5.1, Russel

2020) . with adjustment for multiple testing (Tukey method). Points below the red line indicate significant differences between the seed bank treatments.

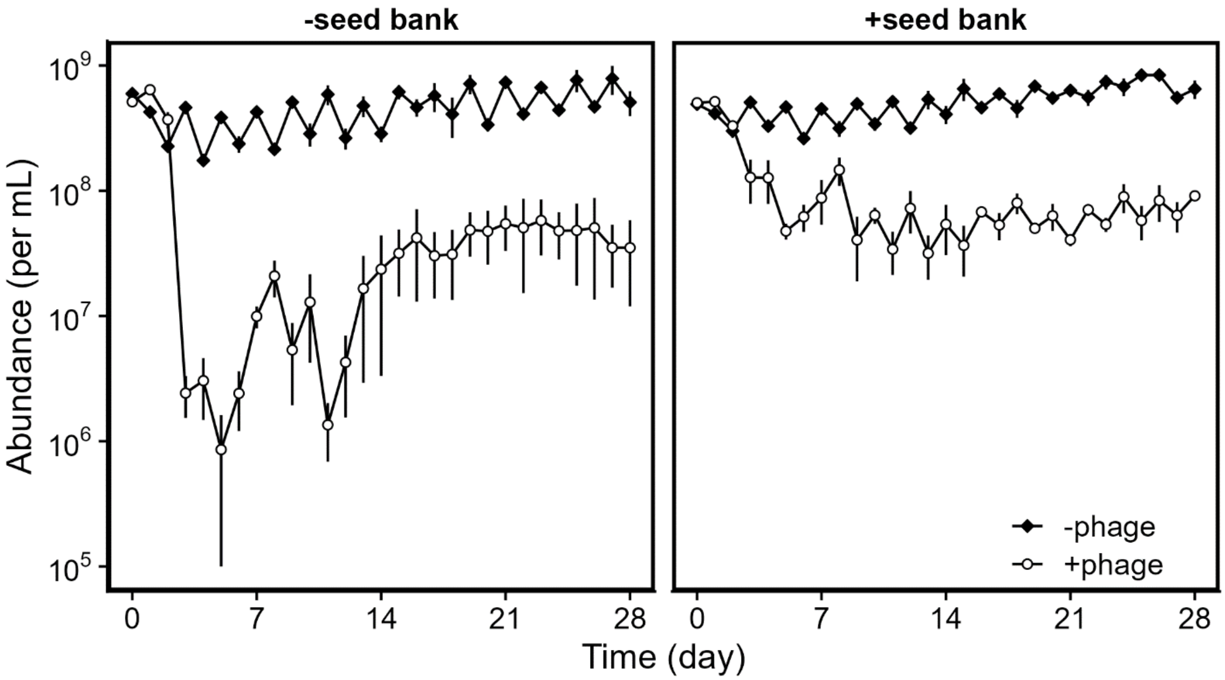


**Fig. S4**. In the absence of phage, host populations (*Bacillus subtilis*) were stable in both seed bank treatments. In the presence of phage, host populations exhibited more pronounced fluctuations, especially in the absence of a seed bank. For all treatments replicate (n = 3) populations were propagated by serial transfer every two days in DSM sporulation media. Data represent mean ± SEM.

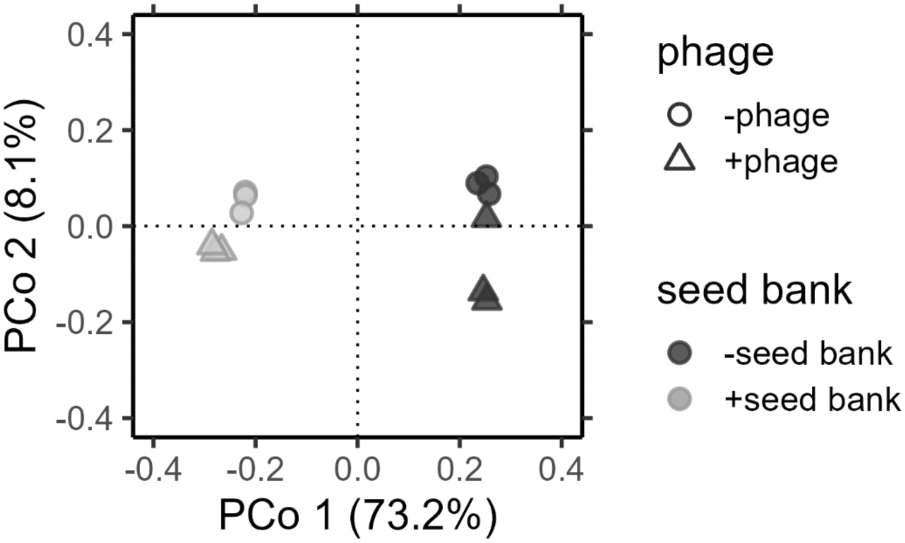


**Fig. S5.** To determine whether host populations in different treatments acquired mutations in different genes, we constructed a gene-by-population matrix in which host genes were scored by the number of mutations acquired at each gene adjusted for gene size and mutation frequency (i.e., *gene multiplicity*). To visualize differences between treatments, we examined the gene-by-population multiplicity matrix using PCoA with a Bray-Curtis distance matrix. Populations (n = 12) separate along PCo1 by seed bank treatment, while populations separate along PCo2 by phage treatment.

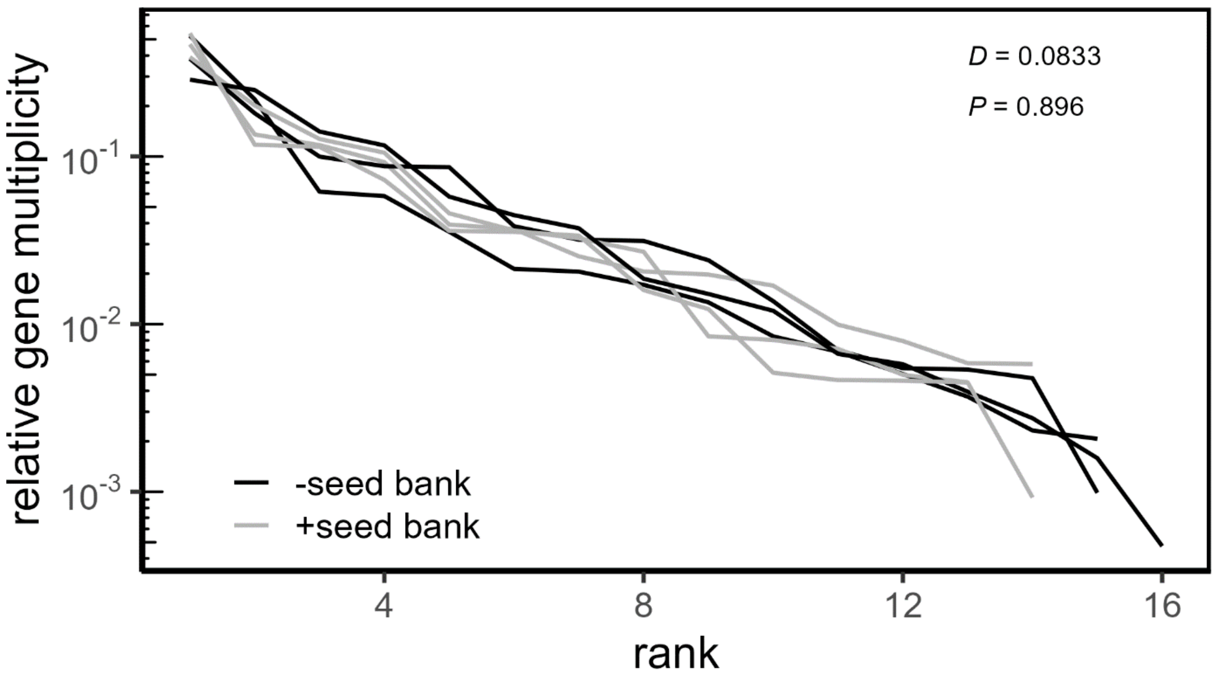


**Fig. S6**. Rank-abundance distribution of acquired mutations across genes of phage SPO1. Gene multiplicity reflects the number of mutations acquired at each gene adjusted for gene size and weighted by the frequency of those mutations in the population. For comparison across populations, we present the relative multiplicity, i.e., the fraction of multiplicity of a gene from the sum of all genes in that population. Each curve shows the relative gene multiplicity ranked by decreasing multiplicity values. The difference between distributions from different seed bank treatments was not significant via a permutational Kolmogorov-Smirnov test.

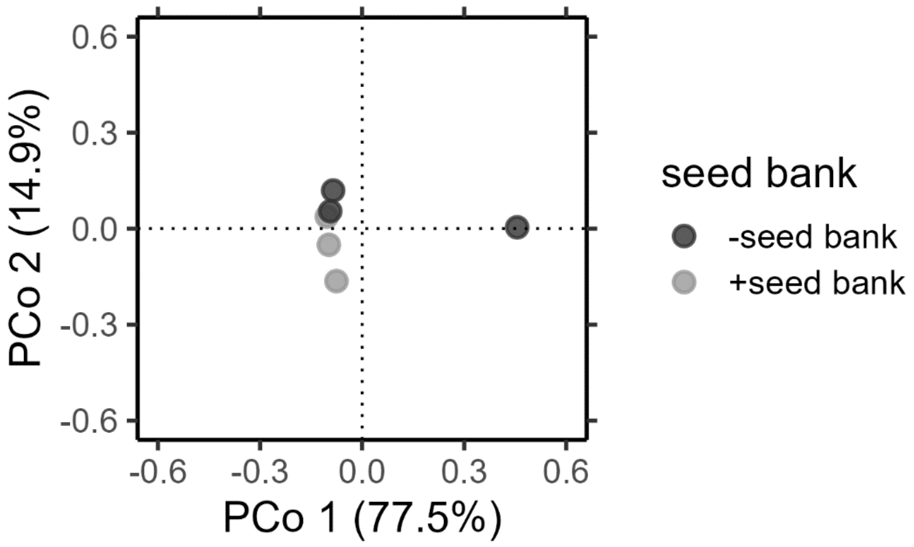


**Fig. S7.** To determine whether phage in different treatments acquired mutations in different genes, we constructed a gene-by-population matrix in which phage genes were scored by the number of mutations acquired at each gene adjusted for gene size and mutation frequency (i.e., *gene multiplicity*). To visualize differences between treatments, we examined the gene-by-population multiplicity matrix using PCoA with a Bray-Curtis distance matrix.

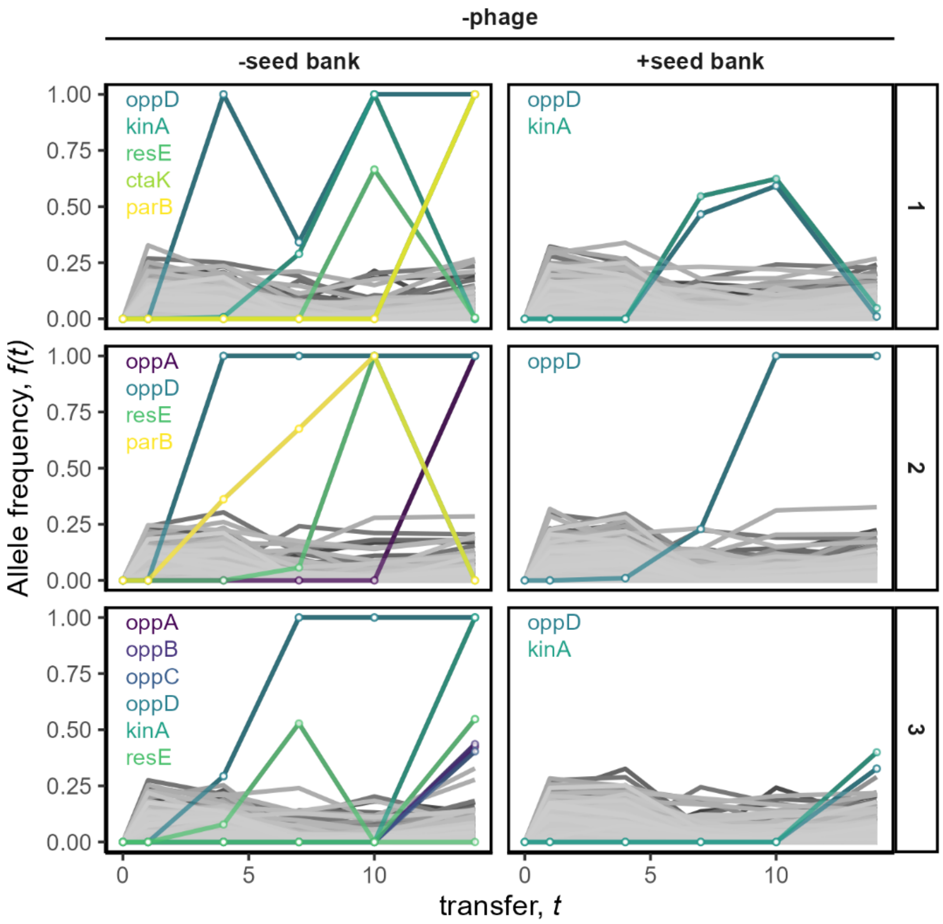


**Fig. S8.** The allele frequency trajectories of host populations evolved in the $-phage$ treatment. Non-synonymous mutations that reached a frequency > 0.3 are colored by the gene represented in the legend. The names of genes with high-frequency mutations are provided for each population. See Table S4 for details on genes.

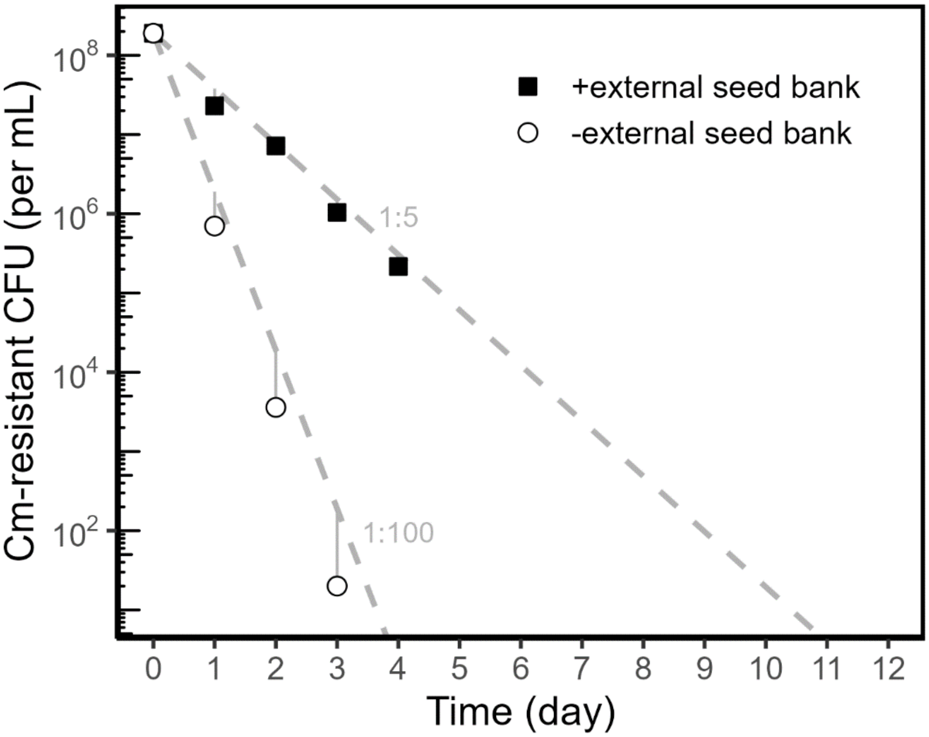


**Fig. S9.** Persistence of spores with and without an external seed bank. Spores derived from a chloramphenicol (Cm) resistant strain of *Bacillus subtilis* that is sensitive to tetracycline were added at day 0 to a serially transferred culture along with a different strain of *B. subtilis* that is sensitive to Cm but resistant to tetracycline, in DSM sporulation medium with tetracycline, so that only the latter strain could grow. The dilution rates expected from the experimental setup are depicted by the grey-dashed lines.

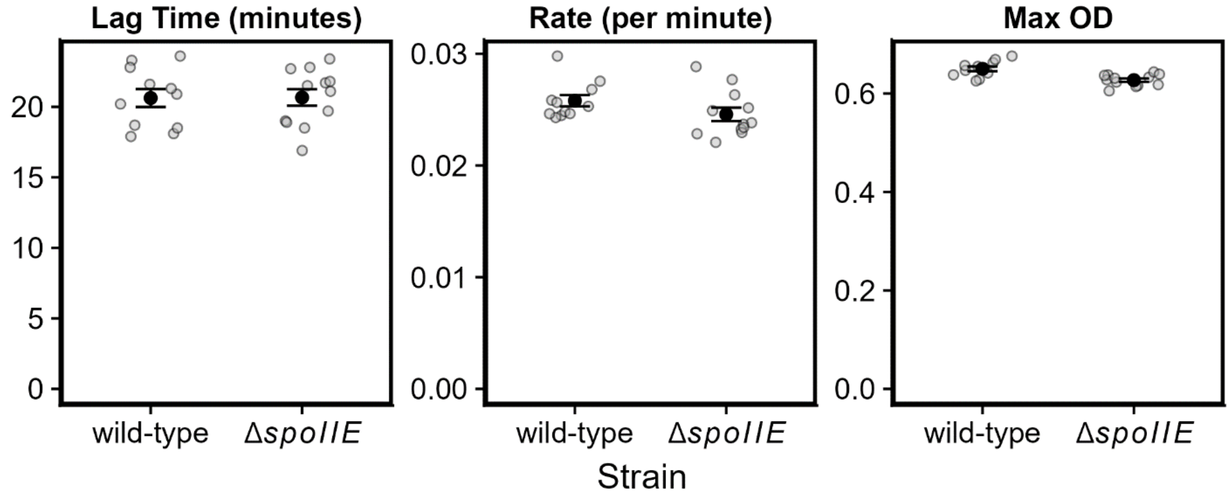


**Figure S10.** Growth parameters for spore-forming (wild-type) and non-spore-forming (Δ*spoIIE*) *Bacillus subtilis* used in the coevolution experiment. In DSM sporulation medium, the two strains have a similar lag time and a similar exponential growth rate. Growth parameters were derived from growth curves (OD600) of three colonies of each strain with four replicate cultures of each (n = 12) using the GrowthRates software (Hall *et al.* 2014). There were no differences between strains in lag time ($t_{20.6 5} = 0.05$, $P = 0.964$) and growth rate ($t_{20.66} = -1.56$, $P = 0.135$) as determined by Welch Two Sample *t*-tests. A difference in max OD was observed ($t_{18.4} = -3.9$, $P = 0.001$).

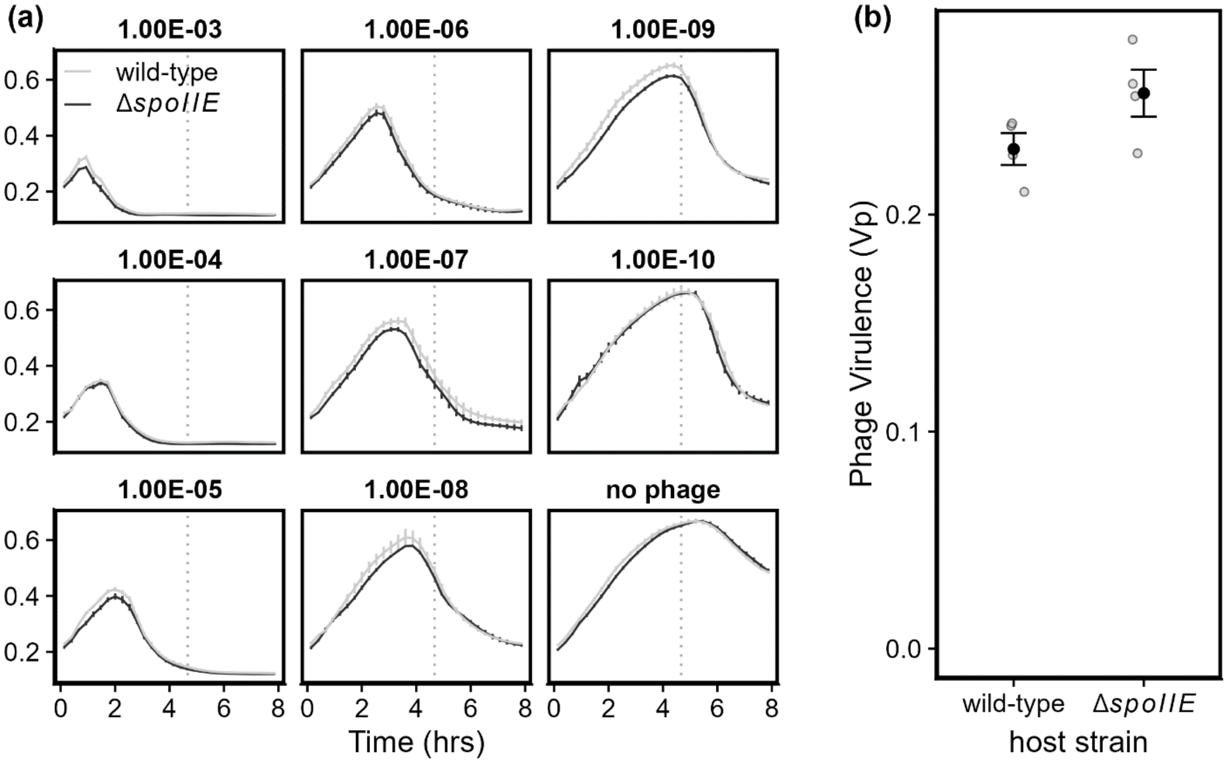


**Fig. S11**. Phage SPO1 has similar virulence towards sporulating *Bacillus subtilis* Δ6 (wild-type) and its non-sporulating derivative (Δ*spoIIE*). Virulence was measured as described by Storms *et al.* (2020). (a) Bacterial density is reduced at the same time and rate across a wide range of infection multiplicities (given in panel titles). Bacterial cultures of each strain were infected with serially diluted phage SPO1. Phage dilution factor is given in panel headers. Curves show mean ± SEM (n = 4). (b) Phage virulence calculated for each host strain from the area under the curves of the data shown in (a) during the exponential growth phase, the end of which is marked by vertical dotted line in (a). Black marks show mean ± SEM of the four individual measurements, shown in grey points. Welch Two Sample *t*-test for virulence on wild-type vs. Δ*spoIIE*: $t_{5.29} = 1.97$, $P = 0.103$.

### SUPLLEMENTARY TABLES

**Table S1.** Bacterial and phage strains used in the study. $Cm^R$ = chloramphenicol resistance; $erm^R$ = erythromycin resistance; $tet^R$ = tetracycline resistance.

| strain | Relevant characteristics | Source (ID) | Ref |
|---|---|---|---|
| *Bacillus subtills* 168 Δ6 | No prophages, $Cm^R$ | BGSC (1A1299) | Westers *et al.* (2003) |
| *Bacillus subtills* 168 Δ6 Δ*spoIIE* | $Spo^-$, No prophages, $Cm^R$ | | This study |
| *Bacillus subtilis* 168 Δ*spoIIE* | *spoIIE*::$erm^R$ | BGSC (BKE00640) | Koo *et al.* (2017) |
| *Bacillus subtilis* 3610 DK7505 | *ganR*::$tet^R$, Blue colonies on X-gal plates | DK7505 | Dempwolff *et al.* (2020) |
| Phage SPO1 | | ATCC (27370-B1) | Stewart *et al.* (2009) |
| Phage SPP1 | Generalized transducing | | Kearns lab, Indiana University |

**Table S2.** Primers used in this study.

| **Name** | **Target** | **Sequence (5’-3’)** | **Source** |
|---|---|---|---|
| oDAS11 | *B. subtilis spoIIE* flanks | TAAGACACCGCCCTTTCACG | This study |
| oDAS12 | *B. subtilis spoIIE* flanks | AGCAGCCATCCGTTATCAGC | This study |
| qSPO1_MCP6.1_F | SPO1 major capsid protein | ACATGAACGCATCTAGCC | This study |
| qSPO1_MCP6.1_R | SPO1 major capsid protein | GCCACCAGTCTGCTTAC | This study |

**Table S3.** *Bacillus subtilis* genes significantly correlated with the first two principal coordinates of the host multiplicity matrix.

*a.* The ten genes (of 87) with the highest positive Pearson coefficient (*r*) significantly correlated with the first principal coordinate axis of the host multiplicity matrix.

| Locus tag (Δ6) | Locus tag (168) | Gene name | Annotation | +/- | *r* | *P*-value |
|---|---|---|---|---|---|---|
| RS08005 | BSU_15370 | *ylmD* | peptidoglycan-editing factor | + | 0.989 | 0.002 |
| RS01655 | BSU_02940 | *yceH* | similar to toxic anion resistance protein | + | 0.979 | 0.002 |
| RS00075 | BSU_00100 | *dacA* | Class C penicillin-binding protein 5 | + | 0.970 | 0.002 |
| RS04460 | BSU_08460 | *yfhA* | ABC transporter for the siderophore schizokinen and arthrobactin (permease) | + | 0.962 | 0.002 |
| RS18745 | BSU_38170 | *qoxA* | cytochrome aa3 quinol oxidase (subunit II) | - | 0.957 | 0.001 |
| RS12620 | BSU_26630 | *yrdQ* | similar to transcriptional regulator LysR | + | 0.953 | 0.001 |
| RS05115 | BSU_09530 | *yhdN* | general stress protein | + | 0.950 | 0.002 |
| RS06930 | BSU_13380 | *ykoS* | unknown | + | 0.940 | 0.001 |
| RS06680 | BSU_12910 | *proG* | 1-pyrroline-5-carboxylate dehydrogenase | + | 0.939 | 0.002 |
| RS05145 | BSU_09600 | *fluC* | flouride exporter | - | 0.938 | 0.002 |

*b.* The ten genes (of 195) with the highest negative Pearson coefficient (*r*) significantly correlated with the first principal coordinate axis of the host multiplicity matrix.

| Locus tag (Δ6) | Locus tag (168) | Gene name | Annotation | +/- | *r* | ***P*-value** |
|---|---|---|---|---|---|---|
| RS01585 | BSU_02800 | *ycdC* | unknown | + | -0.996 | 0.001 |
| RS00185 | BSU_00250 | *xpaC* | hydrolysis of 5-bromo-4-chloroindolyl phosphate | + | -0.995 | 0.001 |
| RS03570 | BSU_06710 | *swrC* | similar to acriflavin resistance protein | + | -0.988 | 0.001 |
| RS19065 | BSU_38780 | *yxkI* | similar to heat shock protein | + | -0.988 | 0.001 |
| RS17980 | BSU_36700 | *moaA* | molybdopterin precursor biosynthesis | - | -0.988 | 0.001 |
| RS14330 | BSU_29890 | *ytnP* | lactonase-homolog protein | - | -0.986 | 0.001 |
| RS11595 | BSU_24090 | *ptb* | phosphate butyryltransferase | - | -0.983 | 0.001 |
| RS10005 | BSU_19230 | *azoR1* | azoreductase | - | -0.978 | 0.001 |
| RS13710 | BSU_28690 | *glcF* | probable glycolate oxidase | + | -0.977 | 0.002 |
| RS17345 | BSU_35490 | *degU* | two-component response regulator | - | -0.976 | 0.002 |

*c.* The genes with positive Pearson coefficient (*r*) significantly correlated with the second principal coordinate axis of the host multiplicity matrix.

| Locus tag (Δ6) | Locus tag (168) | Gene name | Annotation | +/- | *r* | ***P*-value** |
|---|---|---|---|---|---|---|
| RS03905 | BSU_07360 | *yfmS* | soluble chemotaxis receptor | + | 0.774 | 0.006 |
| RS06095 | BSU_11460 | *oppD* | Oligopeptide ABC transporter (ATP-binding protein) | + | 0.695 | 0.019 |
| RS19050 | BSU_38750 | *cydB* | cytochrome bd ubiquinol oxidase (subunit II) | - | 0.661 | 0.028 |
| RS19155 | BSU_38950 | *yxjH* | putative methionine synthase | + | 0.600 | 0.048 |

*d.* The genes with negative Pearson coefficient (*r*) significantly correlated with the second principal coordinate axis of the host multiplicity matrix.

| **Locus tag (Δ6)** | **Locus tag (168)** | **Gene name** | **Annotation** | **+/-** | *r* | ***P*-value** |
|---|---|---|---|---|---|---|
| RS17485 | BSU_35740 | *tagD* | glycerol-3-phosphate cytidylyltransferase | - | -0.785 | 0.009 |
| RS05005 | BSU_09310 | *pgcA* | alpha-phosphoglucomutase | + | -0.701 | 0.007 |
| RS18830 | BSU_38330 | *ywbG* | holin-like auxilary protein | + | -0.611 | 0.025 |
| RS03775 | BSU_07100 | *lplA* | Lipoprotein, putative ABC transporter | + | -0.584 | 0.032 |

**Table S4**. Host and phage genes in which mutations were detected at high-frequency (> 0.3) in serially transferred populations. These genes are highlighted by color and labels in plots depicting mutation trajectories (Figs. 6a, S8).

a. *Bacillus subtilis* genes in which high-frequency mutations were observed. Product and function descriptions are from Subtiwiki (Pedreira *et al.* 2022).

| Name | Product | Function |
|---|---|---|
| *ctaK* | lipoprotein | assembly of the CuA center in Cytochrome caa3 (together with Sco) |
| *gtaB* | UTP-glucose-1-phosphate uridylyltransferase | biosynthesis of teichoic acid |
| *kinA* | two-component sensor kinase | initiation of sporulation |
| *oppA* | oligopeptide ABC transporter (binding protein) | initiation of sporulation, competence development |
| *oppB* | oligopeptide ABC transporter (permease) | initiation of sporulation, competence development |
| *oppC* | oligopeptide ABC transporter (permease) | initiation of sporulation, competence development |
| *oppD* | oligopeptide ABC transporter (ATP-binding protein) | initiation of sporulation, competence development |
| *parB* | CTP hydrolase | chromosome positioning before asymmetric septation |
| *pgcA* | alpha-phosphoglucomutase | interconversion of glucose 6-phosphate and alpha-glucose 1-phosphate |
| *prkA* | AAA+ ATP-dependent protese | control of SigK-dependent gene expression |
| *resE* | two-component sensor kinase | regulation of aerobic and anaerobic respiration |
| *rsiX* | anti-SigX | control of SigX activity |
| *sinR* | transcriptional regulator (Xre family) of post-exponential-phase responses genes | control of biofilm formation |
| *tagD* | glycerol-3-phosphate cytidylyltransferase | biosynthesis of teichoic acid |
| *tagF* | CDP-glycerol:polyglycerol phosphate glycerol-phosphotransferase | biosynthesis of teichoic acid |

*b.* SPO1 phage genes in which high-frequency mutations were observed. Predicted functions are from Stewart *et al.* (2009).

| Gene | Predicted function |
|---|---|
| 1.4 | Unknown |
| 15.1 | Baseplate or tail fiber protein |
| 16.2 | Virion protein (tail?) |
| 18.1 | Tail fiber |
| 18.3 | Tailspike |
| 19.3 | Endolysin |
| 34.32 | Unknown |
| 48 | Unknown |

**Table S5.** Chi-squared goodness-of-fit test results for low-ranking and high-ranking mutated genes compared with proportions of functional categories in the whole genome of the host. See supplementary text for more information.

| rank | seed bank | phage | replicate | statistic | parameters | *P* | $P_{adj}$ |
|---|---|---|---|---|---|---|---|
| low | + | + | 3 | 15.004 | 5 | 0.010 | 0.114 |
| low | + | - | 2 | 13.424 | 5 | 0.020 | 0.114 |
| low | + | + | 2 | 13.261 | 5 | 0.021 | 0.114 |
| low | + | + | 1 | 12.793 | 5 | 0.025 | 0.114 |
| low | + | - | 1 | 10.129 | 5 | 0.072 | 0.240 |
| low | + | - | 3 | 9.842 | 5 | 0.080 | 0.240 |
| high | - | - | 2 | 4.468 | 5 | 0.484 | 0.938 |
| high | - | + | 2 | 4.372 | 5 | 0.497 | 0.938 |
| high | - | + | 1 | 4.208 | 5 | 0.520 | 0.938 |
| high | - | + | 3 | 4.198 | 5 | 0.521 | 0.938 |
| high | - | - | 1 | 3.800 | 5 | 0.579 | 0.947 |
| high | - | - | 3 | 3.198 | 5 | 0.670 | 0.977 |
| high | + | - | 3 | 2.714 | 5 | 0.744 | 0.977 |
| high | + | - | 2 | 2.529 | 5 | 0.772 | 0.977 |
| high | + | + | 2 | 2.185 | 5 | 0.823 | 0.977 |
| high | + | + | 3 | 1.846 | 5 | 0.870 | 0.977 |
| high | + | + | 1 | 0.987 | 5 | 0.964 | 0.977 |
| high | + | - | 1 | 0.804 | 5 | 0.977 | 0.977 |
